# Combined Proteomic and Genetic Interaction Mapping Reveals New RAS Effector Pathways and Susceptibilities

**DOI:** 10.1101/2020.04.20.051623

**Authors:** Marcus R. Kelly, Kaja Kostyrko, Kyuho Han, Nancie Mooney, Edwin E. Jeng, Keene L. Abbott, Dana M. Gwinn, E. Alejandro Sweet-Cordero, Michael C. Bassik, Peter K. Jackson

**Affiliations:** Baxter Laboratory, Department of Microbiology & Immunology, Stanford University School of Medicine, Stanford, California 94305, USA; Program in Cancer Biology, Stanford University School of Medicine, Stanford, California 94305, USA; Division of Hematology and Oncology, Department of Pediatrics, University of California San Francisco, San Francisco, CA 94158, USA; Department of Genetics, Stanford University School of Medicine, Stanford, California 94305, USA; Chemistry, Engineering, and Medicine for Human Health (ChEM-H), Stanford University, Stanford, California, USA; Department of Pathology, Stanford University School of Medicine, Stanford, CA 94305

**Keywords:** KRAS, Pathway Mapping, Interaction Mapping, Synthetic Lethality, Combination Therapy

## Abstract

Activating mutations in RAS GTPases drive one fifth of cancers, but poor understanding of many RAS effectors and regulators, and of the roles of their different paralogs, continues to impede drug development. We developed a multi-stage discovery and screening process to understand RAS function and identify RAS-related susceptibilities in lung adenocarcinoma. Using affinity purification mass spectrometry (AP/MS), we generated a protein-protein interaction map of the RAS pathway containing thousands of interactions. From this network we constructed a CRISPR dual knockout library targeting 119 RAS-related genes that we screened for genetic interactions (GIs). We found important new effectors of RAS-driven cellular functions, RADIL and the GEF RIN1, and over 250 synthetic lethal GIs, including a potent *KRAS*-dependent interaction between *RAP1GDS1* and *RHOA*. Many GIs link specific paralogs within and between gene families. These findings illustrate the power of the multiomic approach to identify synthetic lethal combinations for hitherto undruggable cancers.

**STATEMENT OF SIGNIFICANCE:** We present a thorough survey of protein-protein and genetic interactions in the Ras pathway. These interactions suggested new discoveries that we validate here, and demonstrate important new paralog specificities and redundancies. By comparing synthetic lethal interactions across *KRAS*-dependent and -independent tumors, we identify new combination therapy targets against Ras-driven cancers.

## INTRODUCTION

RAS proteins are mutated in 20% of cancers (1), yet few targeted therapies are available in the clinic (2). Although *RAS* oncogenes were discovered over 35 years ago (3), consequences of RAS activation remain only partly understood. K-, H-, and NRAS are small GTPases that cycle between GTP and GDP-bound states with the aid of exchange factor (GEF) and activating (GAP) proteins. All three are known oncogenes,, but *KRAS* is the most frequently mutated, notably in pancreatic, colon and lung adenocarcinomas (LUAD) (1). Oncogenic mutants of RAS proteins have reduced GAP-mediated GTP hydrolysis, and strongly favor the GTP-bound active state in which they bind tumorigenic effectors. Some RAS effectors can themselves be oncoproteins (2), driving significant interest in them. RAS proteins have many other potentially important effectors and regulators with poorly understood function. Many of these factors are represented by multiple paralogs, which may compensate for one another in screening experiments, concealing their collective importance (4). The extent to which paralogs of RAS effectors or regulators are functionally distinct remains largely unknown. Therapeutic inhibitors against these same proteins, however, might modulate multiple paralogous targets through similar binding sites. The ensemble of RAS protein effectors and regulators thus likely contains new therapeutic targets for unmet clinical needs.

The development of targeted therapies relies on the concept that genetic alterations in cancer cells cause pathway “rewiring,” making nonessential mutant and wild type genes more essential in neoplastic cells (5,6). Targeting such cancer-specific dependencies is a major strategy guiding efforts to treat RAS-driven cancers (2) and has been successful in other cancer types (7,8). Pathway rewiring may also introduce new combination susceptibilities where drugs with functionally linked targets cooperate. While many cancer-specific susceptibilities are known, little systematic investigation of the rewired pathways has been performed. Genomewide knockout screens (9–11) have identified new cancer pathway components, and patient sequencing initiatives (1) have identified driver mutations. Ultimately, however, single gene knockouts and patient genotyping have limited capacity to reveal pathway rewiring because it involves not simply mutation of genes but dysregulation of interactions among them.

Here, we present the integrated results of two direct and orthogonal analyses of interactions in the RAS pathway (Figure 1A). First, we mapped detailed protein-protein interactions (PPIs) using affinity purification/mass spectrometry (AP/MS), which enables unbiased detection of “prey” proteins co-purifying with a chosen “bait”. Beginning with KRAS itself as a bait, we iteratively expanded the PPI network using preys discovered in earlier experiments as bait for later experiments. Second, focusing on the genes identified in this PPI network and on known RAS pathway genes, we mapped genetic interactions (GIs) using CRISPR dual knockout (CDKO) technology (12). CDKO screens identify synergistic or buffering GIs through interference of knockout phenotypes of two genes. GIs link pairs of redundant or interacting genes and suggest new possible combination therapy strategies. From these pairwise links, we constructed networks that produce testable hypotheses regarding the biology of RAS-driven cancers. These results reveal known and new effectors and regulators of the major RAS oncoproteins, show distinct patterns of PPIs between paralogous and non-paralogous proteins, and suggest new connections among RAS and other signaling pathways.

**Figure 1.**
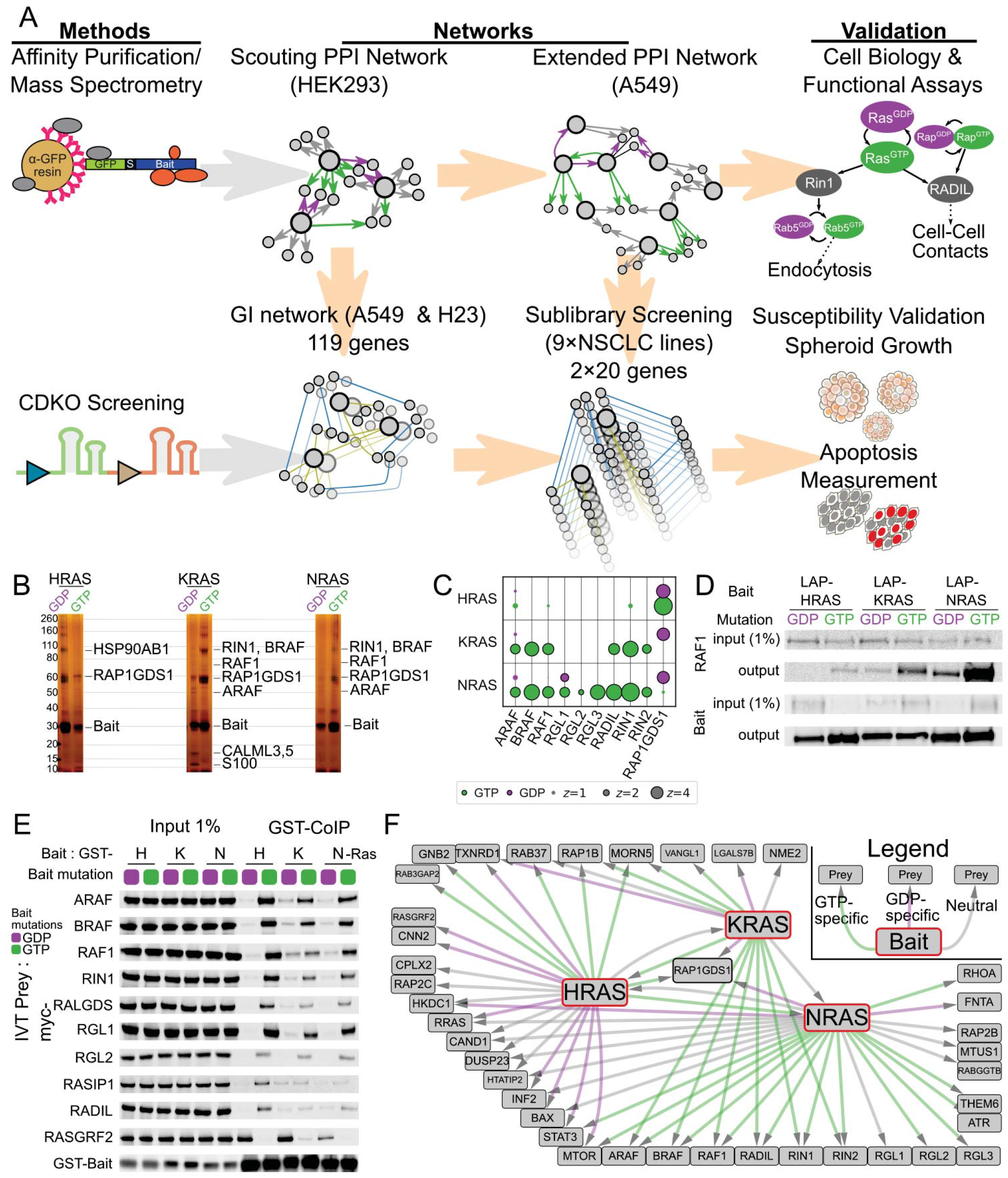
Multiomic Interaction Mapping Reveals Strong Selectivity of RAS Isoforms. **A** Schematic outlining our experimental approach. Top row: Proteins of known interest, expressed as chimeric constructs, were used as bait for AP/MS to discover protein-protein interactions (PPIs). Proteins from the PPI network were selected for CRISPR Dual Knockout (CDKO) genetic interaction (GI) screening. sgRNA cassettes targeting these genes these were transduced into Cas9-expressing *KRAS*-driven LUAD lines and assayed for enrichment or depletion after 21 days growth in culture. Relative depletion of single- and double-targeting cassettes was used to establish synergy or buffering between gene knockout phenotypes. GIs that could form the basis of new therapies were validated by both CDKO and AP/MS in other LUAD lines, as well as functional and growth assays. **B** Silver stain PAGE of tandem affinity purification of H-, K-, and NRAS from A549 cells shows distinct GTP-specific PPIs. Known effectors are marked **C** Bubble plot of 6 AP/MS experiments with GTP- and GDP-locked mutant GTPases as baits (rows), showing the enrichment of selected preys (columns). **D** Western blot of co-immunoprecipitation reactions from A549 cells expressing tagged H, K, or NRAS with mutations locking them in their GTP or GDP-bound forms, probing for endogenous RAF1. HRAS binds significantly less RAF1 than does either other RAS. **E** Cell-free coimmunoprecipitation of selected prey proteins by recombinant purified mutant H, K, and NRAS, imaged by immunoblot. Negligible RAS isoform specificity is observed. **F** Network diagram showing PPIs of RAS proteins and selected proteins. Green and purple lines indicate PPIs from GTP- and GDP-locked preys respectively; gray lines indicate that the PPI is not nucleotide-state specific.

The combined GI and PPI networks connect genes important for LUAD and provide a “wiring diagram” linking drivers of RAS-specific cancer biology. This combination mapping approach provides myriad testable strategies for new combination therapies and is a generalizable method for mapping cell biological pathways altered by disease.

## RESULTS

### Mapping the RAS Pathway through Physical and Genetic Interactions: an outline

Interaction maps were assembled by an iterative process. First, we performed AP/MS of KRAS in HEK293 cells, using previously characterized mutations that confine it to a GTP-locked (2) or GDP-locked (or apo) state (13). Using high-efficiency tandem affinity purification (14,15), we isolated each bait with co-purifying proteins, which were identified by mass spectrometry and quantified as described (16). We expanded the network by repeating the procedure using 5 other RAS pathway proteins as baits, creating a network of 1248 proteins including both known and unknown RAS pathway members (Supplementary Table 2, Figure S1A, S1B).

Guided by this network and using public databases of known cancer driver genes, we selected 119 genes for genetic interaction (GI) analysis by CDKO in the A549 and H23 LUAD cell lines. This analysis of 119 genes (7021 pairs) revealed 999 mostly unknown GIs. We then repeated and expanded the PPI mapping experiments in A549 cells (Supplementary Tables 2 and 3, Figure S2A). Having analyzed the results of both this PPI map and the large GI maps, we created two smaller libraries to screen a set of 9 LUAD lines (see Methods). These cell lines show a range of dependency on *KRAS*, allowing us to stratify GIs into classes more or less dependent on KRAS protein activity. Here we describe new insights from these networks into *KRAS*-driven cell biology and potential combinatorial susceptibilities (Figure 1A, S1C).

### Affinity Purification/Mass Spectrometry Mapping Reveals Strong Selectivity of RAS Isoforms

Activated, oncogenic forms of Ras proteins cause transformation in specific tumors through enhanced binding to downstream effectors (2). Many effectors bind one or more RAS isoforms, but no systematic analysis of RAS PPIs with their regulators and effectors has been performed. From AP/MS experiments in HEK293 cells, we identified interactors of KRAS, both well-known (RAFs) and less studied (RALGEFs, RAP1GDS1, RADIL) (Figure S1B). As a comparison, we conducted copurifications with GTP- and GDP-locked mutants of H-, K-, and NRAS in A549 cells, a *KRAS*-transformed LUAD line (Figure S2A, S2B).

Differences in co-purifying proteins between each RAS protein, and between their mutant forms, were apparent by silver stain (Figure 1B) and by mass spectrometry (Figure 1C). RAS effector proteins with known preference for the GTP-locked state bound specifically to that mutant form. Surprisingly, there were notable differences among the specific effectors bound by activated H-, K-, and NRAS (Figure 1C).

Most strikingly, RAF kinases (ARAF, BRAF, RAF1) were highly enriched with either GTP-locked K- or NRAS baits but not HRAS. We analyzed small-scale coimmunoprecipitations in lung cancer cells by immunoblot (Figure 1D) and confirmed that RAF1 bound GTP-locked HRAS much less efficiently than GTP-locked K- or NRAS, even with similar expression of RAS bait proteins. The same was true for three other RAS effectors containing canonical RAS-association (RA) domains: RIN1, RIN2, and RADIL. In contrast, three members of the RALGEF family – RGL1, −2, and −3 – that also contain RA domains copurified efficiently with NRAS but poorly with H- or KRAS.

We considered whether observed binding specificities reflected protein-intrinsic properties that could be recapitulated in cell-free assays. Using purified GTP- and GDP-locked H-, K-, and NRAS proteins, we reconstituted binding to *in vitro*-translated forms of the known and new RAS-interacting proteins identified (Figure 1E, S2C). Although effector PPIs were strongly GTP-dependent, we found negligible differences in binding efficiency between the three GTP-locked RAS proteins and these RAS effector proteins *in vitro*. This and experiments below indicate that other cellular processes (such as regulation of localization) are comparably important to RAS nucleotide state in determining RAS-effector engagement in tumor cells.

A number of known RAS-binding proteins were not identified by AP/MS. PPIs between GTPases and their coordinate GEFs have been historically difficult to isolate by AP/MS (see (17)), so the absence of signal from GEFs like SOS in these purifications was expected. The absence of other RAS-binding proteins, like PIK(3)-CA or TIAM1, requires further study. These PPIs may only occur in specific cell lines or under stimuli not recapitulated in our culture method.

Combining this AP/MS data with PPI data from BioGRID (18), we constructed a PPI network of RAS interactors (Figure S2A), a selective representation of which is shown in Figure 1F. Though these PPIs are efficiently observed by AP/MS, not all are necessarily direct (see (14)). Many PPIs suggest functional clues for how the RAS-effector PPIs that we observe are regulated.

### AP/MS Identifies New KRAS Effectors Underlying RAS-Driven Macropinocytosis and Migration

Interactions between RAS family proteins and effectors are mediated by the effector binding domain of RAS proteins and RA domains in target proteins (19). In addition to the effectors of KRAS mentioned above, we found two proteins that co-purified efficiently with both GTP-locked K- and NRAS that contain canonical RA domains (Figure 2A): RIN1 and RADIL.

**Figure 2.**
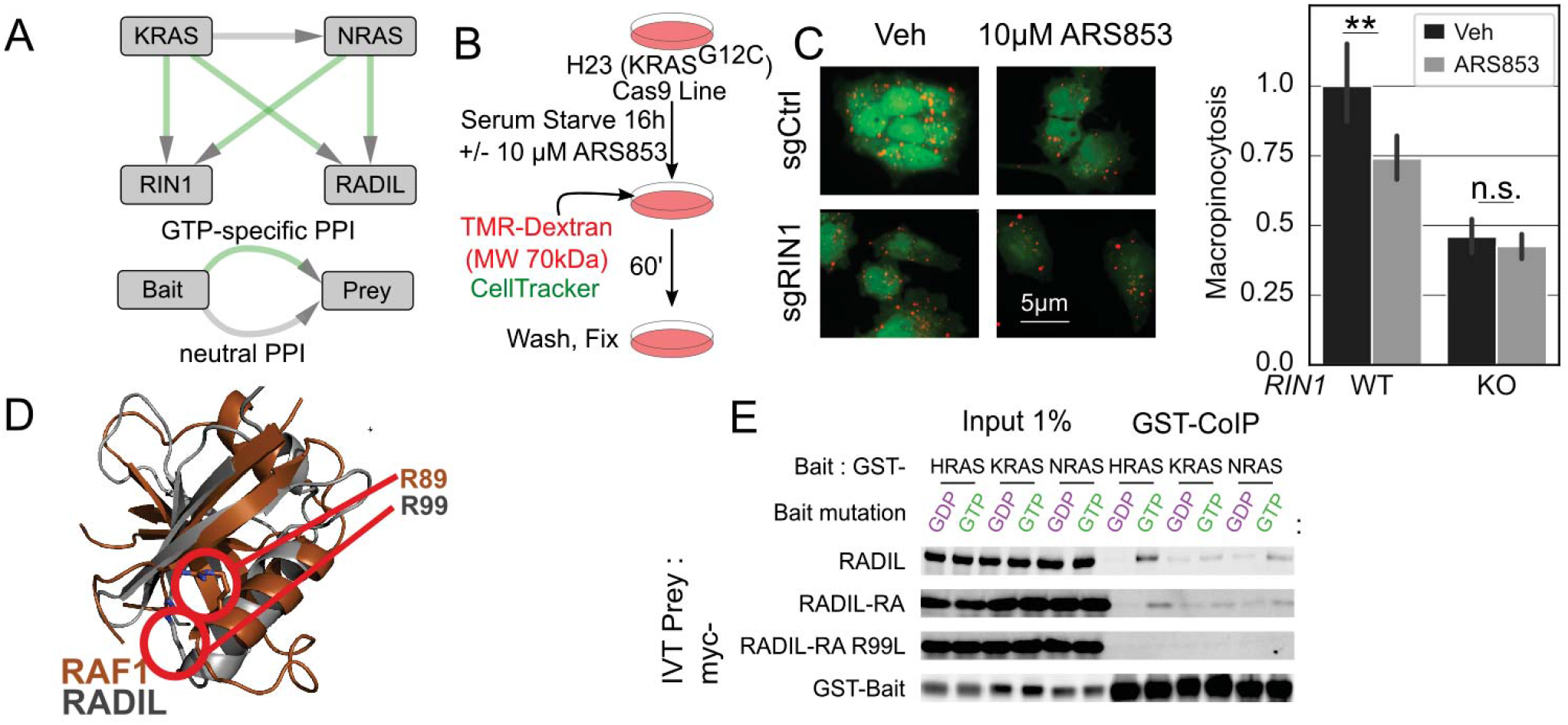
AP/MS Identifies New KRAS Effectors Underlying RAS-Driven Macropinocytosis and Migration. **A** Network diagram showing PPIs between RAS proteins, RIN1 and RADIL. **B** Schematic of assay to assess macropinocytosis (MPC), derived from (24). **C** MPC was quantified by TMR-dextran (red) signal under fluorescence microscopy. RAS inhibition reduced dextran uptake in wild-type, but not RIN1 knockout cells. Error bars indicate 95% confidence intervals estimated from 76 fields from the wild type and 226 from knockouts (Supplementary Methods) **D** Left: Superposition of crystal structures of the RA domain of RAF1 (PDB ID: 4G0N, brown) and RADIL (PDB ID: 3EC8, gray). RADIL R99 superimposes near RAF1 R89, which is required for RAS-RAF binding (19). **E** Cell-free co-immunoprecipitations indicated myc-tagged constructs as prey, and indicated GST-RAS fusion proteins as bait. myc-RADIL binds RAS directly, and its RA domain (residues 61-164) is sufficient. R99L is required.

RIN1 was previously shown to bind HRAS *in vitro* (20), and functions as a nucleotide exchange factor for RAB5 GTPases, which are central organizers of endocytosis (21). Recent work shows *KRAS*-transformed cancers use macropinocytosis, a specific form of endocytosis, to scavenge nutrients from their environment (22,23). The observation of RAS-RIN PPIs suggested that KRAS might directly regulate macropinocytosis through RIN1.

We assayed macropinocytosis as described (24) (Figure 2B). Serum-starved H23 cells take up 70kDa fluorescent dextran by macropinocytosis, as visualized by fluorescence microscopy (Figure 2B). Three clonal CRISPR *RIN1* knockout lines (Figure S3A) showed decreased macropinocytosis relative to cells transduced with a negative control sgRNA. Upon pretreatment with ARS853, a KRAS^G12C^-specific inhibitor (25), macropinocytosis was substantially suppressed in the control cells but not in the *RIN1* knockout cells. These cells appear insensitive to further inhibition of macropinocytosis by ARS853, consistent with the model that RAS-driven macropinocytosis is mediated largely through RIN1 (Figure 2C).

RADIL also strongly co-purified specifically with GTP-locked KRAS in both A549 and HEK293 lines. The N-terminal RA domain in RADIL resembles that of other RAS effectors (Figure S3B) and has been crystallized (26). Arginine 99 (R99) in RADIL occupies a position in the RA domain comparable to RAF1 R89, a site required for RAS interaction (19) (Figure 2D). We find that RADIL specifically and efficiently binds *in vitro* to GTP-locked H-, K-, and NRAS, that the RA domain is sufficient for this PPI and that the R99 mutant fails to bind KRAS-GTP (Figure 2E, S3C). Previous studies of RADIL focus on its role in RAP1A-driven cell adhesion (27), and our results suggest a direct link between KRAS and RADIL, at least in lung cancer. To validate that RADIL also affects adhesion and motility in these cells, we conducted a scratch-wound assay (28) and found that RADIL knockdown increased the migration of these cells (Figure S3E-G), in agreement with other studies (27).

Below, we interrogate genetic interactions (GIs) between RAS pathway components, including RIN1 and RADIL. Though we observe synthetic lethal GIs involving both effectors, they are fewer than those involving other effectors like RAF kinases. Given the importance of macropinocytosis for scavenging nutrients, we expect that RIN1 would be more significant in more nutrient-poor conditions than in rich media. Similarly, given its role in migration, RADIL may assume more significance in tumor metastasis than growth control.

### Systematic Mapping of the RAS/RAP/RAL Signaling Pathway Shows a Network of Interconnected GTPases

We identified strong GTP-dependent PPIs between KRAS and the RALGEFs RALGDS, RGL1, and RGL2 in HEK293 cells. While RALGEFs, and their RAL GTPase substrates, are linked to cell proliferation and carcinogenesis (29–31), they remain incompletely understood. RALGEFs also bind RAP GTPases and different family members have distinct but unknown RAL-independent activities (31,32).

We conducted AP/MS experiments with RALGDS, RGL1, and RGL2 as bait proteins. Each RALGEF co-purified a variety of upstream RAS and RAP GTPases (Figure 3A). We identified other interactors unique to each RALGEF, implicating them in signaling pathways including the Hippo pathway (NF2) and NF-κB (NKIRAS2).

**Figure 3.**
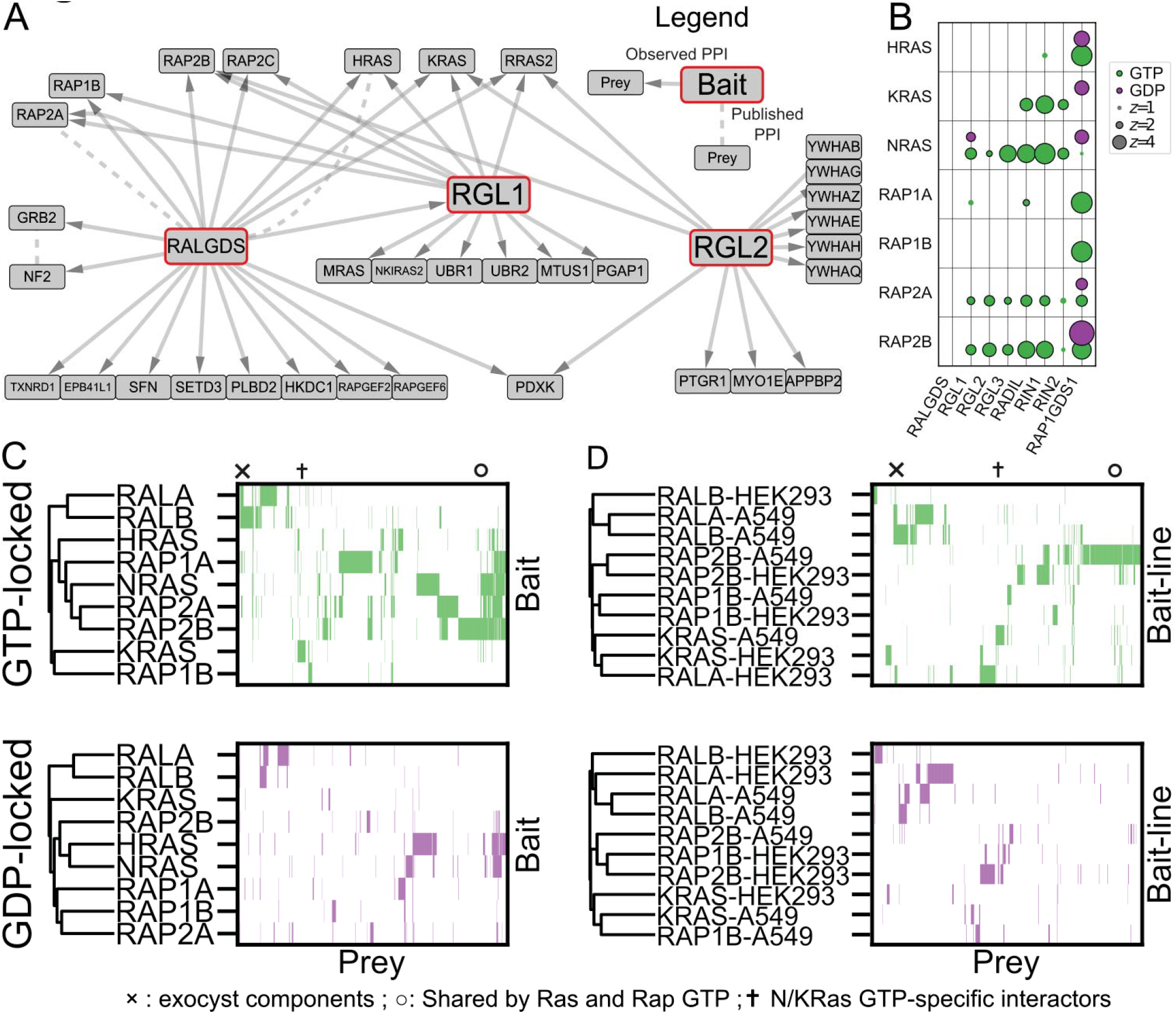
Systematic Mapping of the RAS/RAP/RAL Signaling Pathway Shows a Network of Interconnected GTPases. **A** Network diagram of PPIs in A549 cells with RALGEF family members as baits. The RALGEFs differ in their PPIs. **B** Bubble plot of 14 AP/MS experiments with GTP- and GDP-locked mutant GTPases as baits (rows), showing the enrichment of selected preys (columns). RAP2A and RAP2B bind some canonical RAS effectors in their GTP-locked states. **C** Clustermap of prey proteins significantly enriched (FDR<0.05) in each experiment with the indicated baits in A549 cells. Prey protein order is preserved across panels C and D, but prey proteins not significantly enriched in at least one experiment were removed for simplicity. Symbols indicate columns in the clustermap corresponding to proteins of interest: × Exocyst components; † N/KRAS-GTP specific interactors; ∘ Common interactors of K- and NRAS and of RAP2. **D** Clustermaps as in C but comparing GTPases that were used as bait proteins for AP/MS experiments in both A549 and HEK293 cells.

To better understand RALGEF interaction with specific GTPases, we conducted AP/MS experiments with the RAP and RAL GTPases in their GTP- and GDP-locked forms. RAP proteins engage effectors, including the RALGDS family, through effector RA domains. A comparison of common PPI partners of RAS and RAP family members in A549 cells is shown in Figure 3B, indicating that RADIL, RALGEF and RIN-family effectors are preferentially engaged by NRAS over H- or KRAS, and RAP2A and -B over RAP1.

Clustering analysis of RAS, RAP, and RAL GTPases showed that RAS and RAP GTPases bind surprisingly similar effectors including RALGEFS, RIN1/2 and RADIL (marked ∘). In contrast, the RALA and -B GTPases show a distinct set of interactors (Figure 3C) including all components of the exocyst complex (marked ×), which controls exocytosis. A subset of AP/MS experiments were conducted in both A549 and HEK293 cells. An analogous clustering analysis (Figure 3D) shows that, generally, each GTPase interacted with a similar set of proteins in both lines. Most strikingly, clustering analysis does not distinguish RAS from RAP proteins, indicating an underappreciated similarity between RAS and RAP signaling.

These PPI maps show the benefit of systematic mapping of paralogs and similar families. The many specific PPIs of RAP, RAL, and RALGEF family proteins suggest the existence of a network of dynamic interactions controlling signaling and trafficking at plasma and organellar membranes. A curated subset of these PPIs is diagrammed in Figures S1A and S2A. Complete PPI data can be found in Supplementary Table 2, and <u>online via NDEX</u>. Importantly, these PPIs predict and rationalize many genetic interactions, as we show below.

### CRISPR Dual Knockout Screens of KRAS Interactome Components Reveal Functional Relations and Potential Susceptibilities

Having established that this high-confidence PPI network contains many uncharacterized but potentially important interactors, we next sought to map genetic interactions (GIs) between them to better understand the RAS pathway and identify combination target pairs.

We constructed a sgRNA library targeting 223 genes based on their protein identification in AP/MS experiments (Figure S1B, Figure S4A, B, Supplementary Table 4). The library contained 10 sgRNAs per gene and was transduced into A549-Cas9 cells. Cells were grown for 21 days and sequenced to identify depleted or enriched sgRNAs. This screen identified some essential genes, which would be less likely to show GIs and so were removed from further consideration. Using a combination of proteomic and genetic criteria, we selected 109 genes from the network for CDKO screening. To these we added genes encoding MEK and ERK kinases, and others reflecting genetic lesions in A549 cells for a total of 119 genes (Figure 4A, Supplementary Table 5).

**Figure 4.**
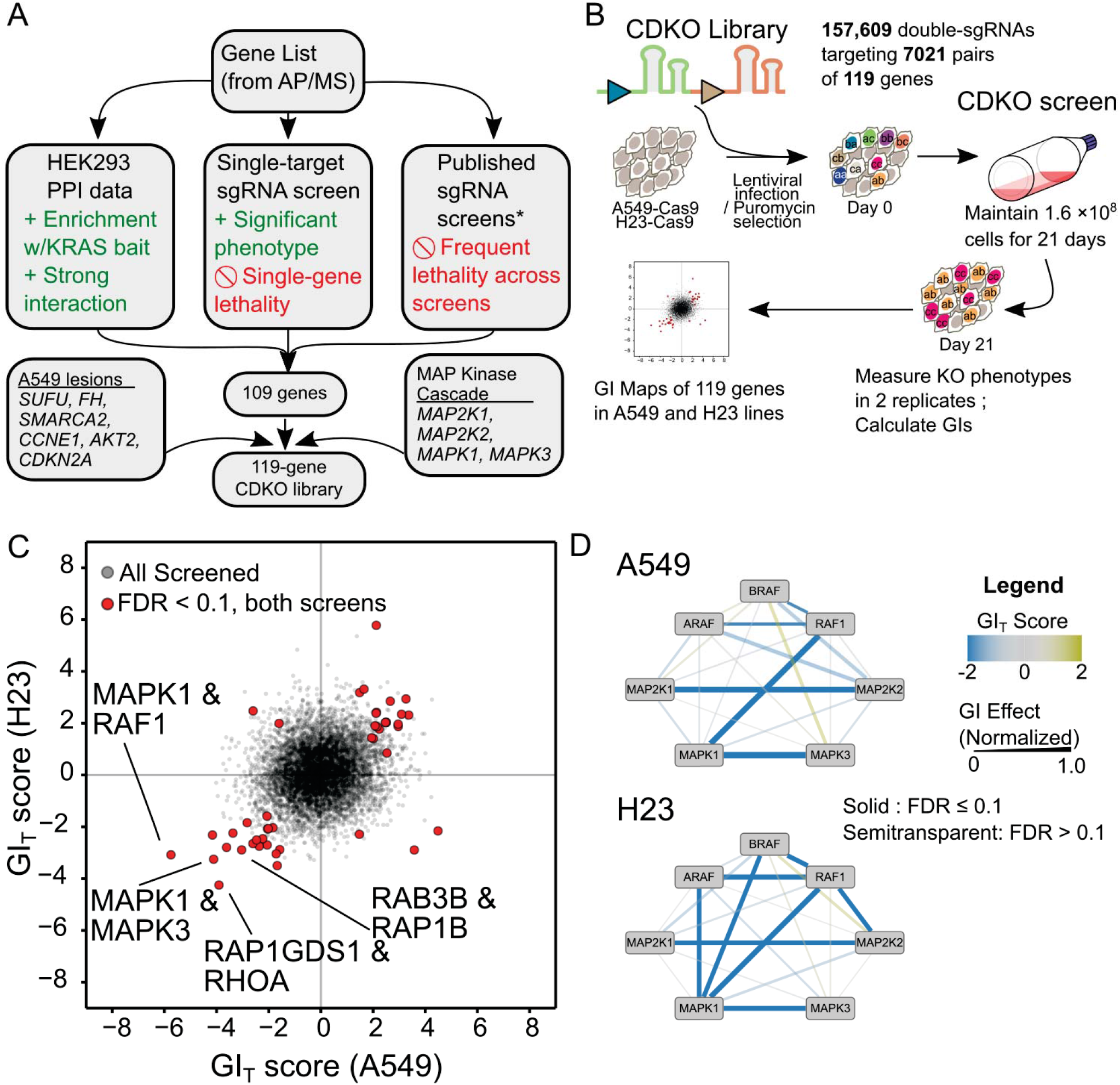
CRISPR Dual Knockout Screens of KRAS Interactome Components Reveal Functional Relations and Potential Susceptibilities. **A** Schematic illustrating gene selection for the CDKO (CRISPR Dual Knockout) screen. Genes involved in strong PPIs, especially with KRAS, were prioritized, while genes producing weak or outright lethal phenotypes were discarded. **B** Schematic showing CDKO screening. Cas9-expressing LUAD cells are transduced with pairs of sgRNAs. Cells are sequenced before and 21 days after transduction. Relative sgRNA abundances are used to calculate their effects. **C** Scatterplot showing GIs in the A549 and H23 lines. Negative, “synergistic,” phenotypes are more severe than would be expected from the two single-gene effects in combination. **D** GIs between components of the MAPK pathway in A549 (left) cells and H23 (right) cells differ.

To measure GIs among these genes, we constructed a CDKO library including 3 sgRNAs per gene (12) to enable deletion of all 7,021 pairs. We included 60 “safe” sgRNAs targeting nonfunctional genomic loci to control for intrinsic effects of Cas9 activity and allow measurement of single gene deletion phenotypes. The library was transduced into two Cas9-expressing KRAS-mutant LUAD lines, A549 and H23. The abundance of each guide pair was measured before and after the 21-day growth assay (Figure 4B). By comparing the abundance of gene/safe guide pairs and double-targeting guide pairs, GIs were quantified (Figure S4C,D, Supplementary Table 5, Methods).

We identified 548 significant GIs in A549 cells, and 447 in H23 cells, of which 59 overlapped (Figure 4C). The consistently interacting pairs include paralogs such as *RAP1A* and *RAP2A* or genes in the same pathway, such as *MAPK1* and *RAF1*. Many of the strongest synergistic GIs in either line were found between paralogous MAPK pathway components (e.g. *MAPK1* and *MAPK3, MAP2K2* and *MAP2K4*). These strong dependencies on this known central pathway confirms that the screening captured relevant GIs. By combining GI and PPI data, we found that roughly one in six GIs paralleled direct (first-degree) PPIs, while many more GI partner pairs are connected to second or third-degree PPIs (Figure S5A).

Representative GI/PPI subnetworks, showing GIs common to both cell lines tested, are diagrammed in Figure S5B. These combined maps provide testable hypotheses about mechanisms underlying observed GIs. For example, we observe a synergistic GI between a PI(3) kinase subunit *PIK3R1* and *RAF1*. Both represent RAS-related pathways, and this GI is supported by pharmacological observations (33).

GI strength is quantified by GI_T_ score, which describes the degree of synergy of a given GI, and accounts for assay noise (12). As one strong example, KRAS-transformed H23 cells infected with the safe/safe, *RAP1GDS1*/safe, and safe/*RHOA* sgRNA cassettes doubled once every 51, 53 and 64 hours, respectively (Figure S8A). Those infected with a double-targeting guide doubled only every 120 hours. The efficacy of this combination is paralleled in multiple assay systems, as we show below. These changes in doubling time correspond to a GI_T_ score of −4.2. Strong GI_T_ scores may therefore indicate dramatic changes in doubling time.

We included MAPK pathway genes *MAP2K1, MAP2K2, MAPK1*, and *MAPK3* in the CDKO library as positive controls because of known synthetic lethal GIs among these genes (34). As predicted, we observed strong synthetic lethal GIs between these paralogous pairs. Combined knockout of *MAPK1* and *RAF1* was also strongly synergistic, but surprisingly, knockout of *RAF1* and *MAPK3* was much less so. A549 cells showed strong dependency on *RAF1* in combination with other pathway members, but limited dependency on *ARAF* or *BRAF*. In H23 cells, however, GIs could be observed between all three *RAF* genes and other MAPK pathway components (Figure 4D). The seven MAPK genes showed numerous but highly varied GIs with other genes in the two cell lines (Figure S5C). This suggests that the MAPK pathway signals to other RAS effector pathways, but in different combinations in different tumor lines.

Though the presence of synergistic GIs common to both lines indicated that CDKO screening could uncover important combination susceptibilities, the preponderance of cell line-specific GIs indicated screening more cell lines to discern consistent patterns of susceptibility. In particular, we wished to determine which synthetic lethal GIs were specific to *KRAS*-dependent cancers.

### GI Mapping Identifies Exploitable Synergistic Interactions in RAS-Driven Lung Adenocarcinoma

To identify GIs specific to *KRAS*-driven LUAD, we selected nine lines, deleted *KRAS* in those cells using CRISPR/Cas9, and measured their growth to determine their *KRAS*-dependence. We found that six cell lines were *KRAS*-dependent, and three cell lines were *KRAS*-independent (Figure 5A, S6A-C). Therapeutically useful combination target pairs should show synergistic lethal GIs in *KRAS*-dependent, but not *KRAS*-independent lines. We constructed two smaller CDKO libraries to conduct focused screens across the cell line panel. The first, “GI-Directed” library, targeted 20 genes with strong and numerous GIs identified in the first GI screen. The second, “paralog” library, targeted paralogous families of RAS regulator and effector proteins, focusing on those found in the A549 PPI network (Supplementary Tables 6 and 7).

**Figure 5.**
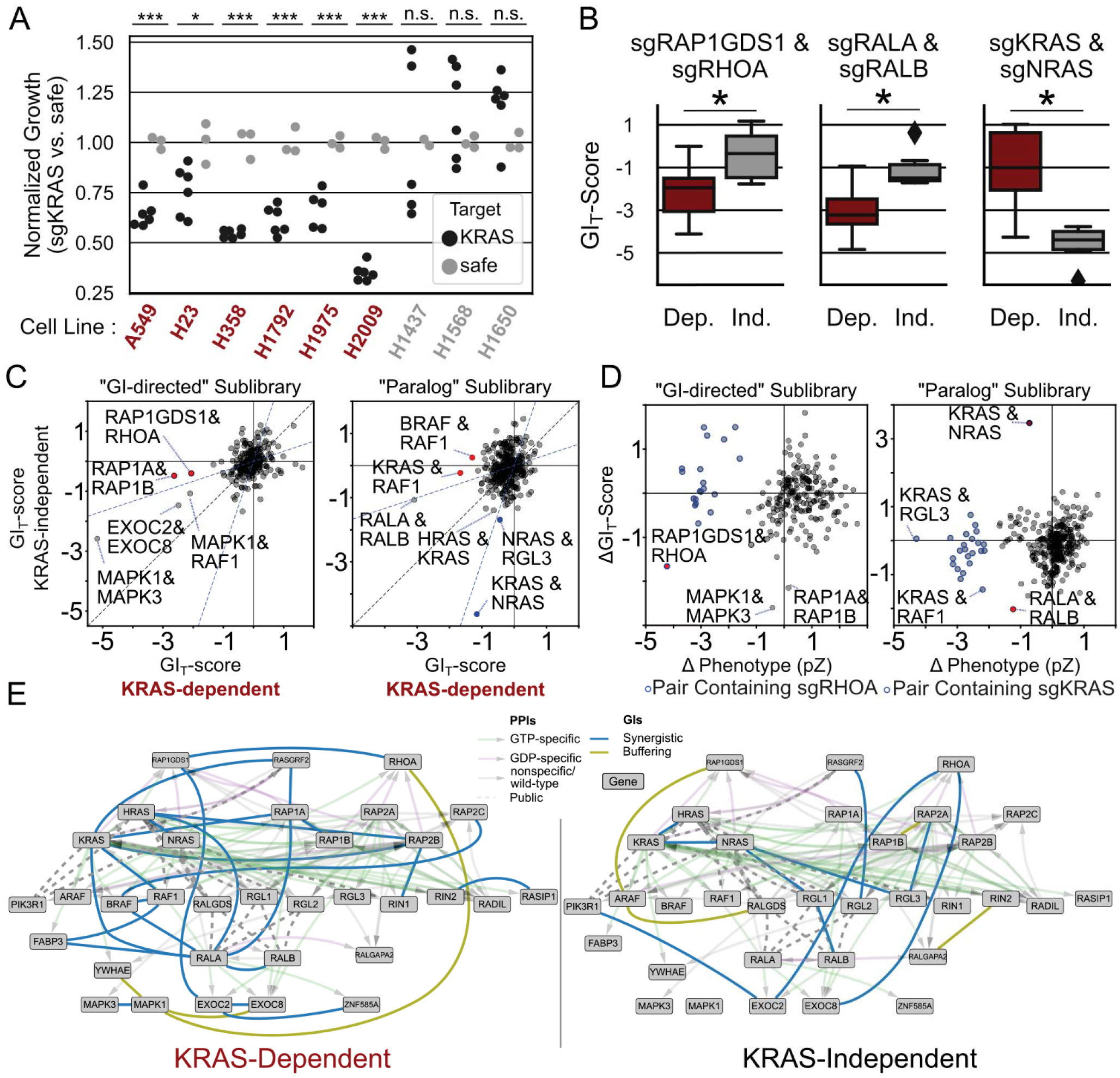
GI Mapping Identifies Exploitable Synergistic Interactions in RAS-Driven Lung Adenocarcinoma. **A** Nine engineered LUAD lines stably expressing Cas9 were assayed for *KRAS* dependence. Lines were stratified into *KRAS*-dependent (labeled red) and *KRAS*-independent (gray) cohorts. * *p* < 0.05 *** p < 0.001 by one-sided *T* test. **B** Two smaller “sublibraries” were screened for GIs in the lines identified in panel A. One sublibrary (“GI-Directed”) targeted the strongest interactors from the first screens, and the other (“paralog”) was targeted families of RAS-interacting proteins. GIs were identified with different patterns of interaction in the *KRAS*-dependent (red; *n*=12) or –independent (gray; *n*=6) cohorts. The GI_T_ scores of each GI across individual replicates are rendered as box plots. *: *p* < 0.05 by Mann-Whitney U test. **C** Scatter plot of mean GI_T_ score for each gene pair in *KRAS*-dependent (*x* axis) and –independent cohorts (*y* axis) in both sublibraries. Noteworthy gene pairs labeled. **D** Scatter plot of difference in observed double-KO phenotype (*x* axis) vs. mean GI_T_ score (*y* axis). The average GI_T_-score for each gene pair across each set of lines is plotted for each library. The dashed line indicates equal phenotypes in both sets. The *RAP1GDS1/RHOA* pair is uniquely potent and toxic in *KRAS*-dependent lung cancer lines. **E** Combined PPI/GI networks. GIs enriched to one cohort or another (determined by Mann-Whitney U Test) are plotted in each network.

In both screens we found examples of gene pairs whose synergy was greater in the *KRAS*-dependent than -independent lines. Among these pairs was *RAP1GDS1* and *RHOA* (Figure 5B, left), which had shown a strong synthetic lethal GI when screened in the earlier, larger libraries. Another strong synthetic lethal pair specific to *KRAS*-dependent lines was *RALA*/*RALB* (Figure 5B, middle), which previously demonstrated synthetic lethality in genetically engineered mouse models (34). In contrast, *KRAS* and *NRAS* double knockout strongly synergized specifically in *KRAS*-independent lines (Figure 5B, right). *KRAS*-dependent cells may have become “addicted” specifically to *KRAS*, but *KRAS*-independent cells may rely on K- and NRAS proteins for similar signaling functions.

Many other GIs common to both A549 and H23 cells in the original screen, such as *MAPK1/MAPK3*, and *MAPK3/RAF1*, were recapitulated in the sublibrary screens. GI_T_ scores of screened gene pairs are plotted in Figures 5C and D. Many had a greater tendency toward genetic interaction in either *KRAS*-dependent or -independent lines. Among these GIs, the *RAP1GDS1/RHOA* pair had both uniquely strong synergy and toxicity in *KRAS*-dependent lines (Figure 5C). In the GI-directed sublibrary, *KRAS*-dependent lines were found to have many specific genetic GIs involving *RHOA* (blue circle outlines) (Figure 5D).

GIs specific to either the *KRAS*-dependent or -independent cohort are plotted over semitransparent PPI networks in Figure 5E. While few synthetic lethal pairs involving one MAPK pathway component were consistently more severe across the *KRAS*-dependent lines, there were significantly more such GIs in the *KRAS*-dependent than -independent cohorts (Figure S6D). All GIs observed in all 9 cell lines and two libraries are diagrammed in Supplementary Figure S7.

### *RAP1GDS1* and *RHOA* form a Synthetic Lethal Gene Pair in RAS-Driven Lung Adenocarcinoma

As mentioned, we observed a synthetic lethal GI between *RAP1GDS1* and *RHOA* that was significantly stronger in *KRAS*-dependent cells. *RHOA* encodes a small GTPase critical for actin assembly and membrane organization (35). RAP1GDS1, or SmgGDS, is less well characterized but has serves as GEF for RHOA and RHOC (36). RAP1GDS1 also binds other small GTPases including KRAS (37). RAP1GDS1 differs structurally from canonical RHO and RAS GEFs (38) and may also act as a chaperone for these small GTPases (Berg et al, 2010; Shimizu et al, 2018; Schuld et al, 2014). RAS-driven oncogenesis also depends on RHO proteins (39). The CDKO screens point to specific GIs among *RAP1GDS1*, *RHOA* and mutant *KRAS* not previously reported.

To validate the *KRAS*-specificity of this GI, we used the CRISPR/Cas9 system to knockout *RHOA, RAP1GDS1*, or both genes together in a set of *KRAS*-dependent and – independent LUAD cell lines and in human bronchial epithelial cells (HBECs) (Figure S8A). Double *RAP1GDS1* and *RHOA* knockout selectively impaired proliferation of *KRAS*-dependent LUAD lines but had minimal synergy in *KRAS*-independent lines (Figure 6A, S8B). Neither single-gene knockout had a strong effect on proliferation in these cells, suggesting that only the loss of both genes confers a vulnerability in *KRAS*-dependent cells. Analysis of apoptosis by flow cytometry revealed that double knockout induced cell death only in *KRAS*-dependent cells (Figure 6B, C).

**Figure 6.**
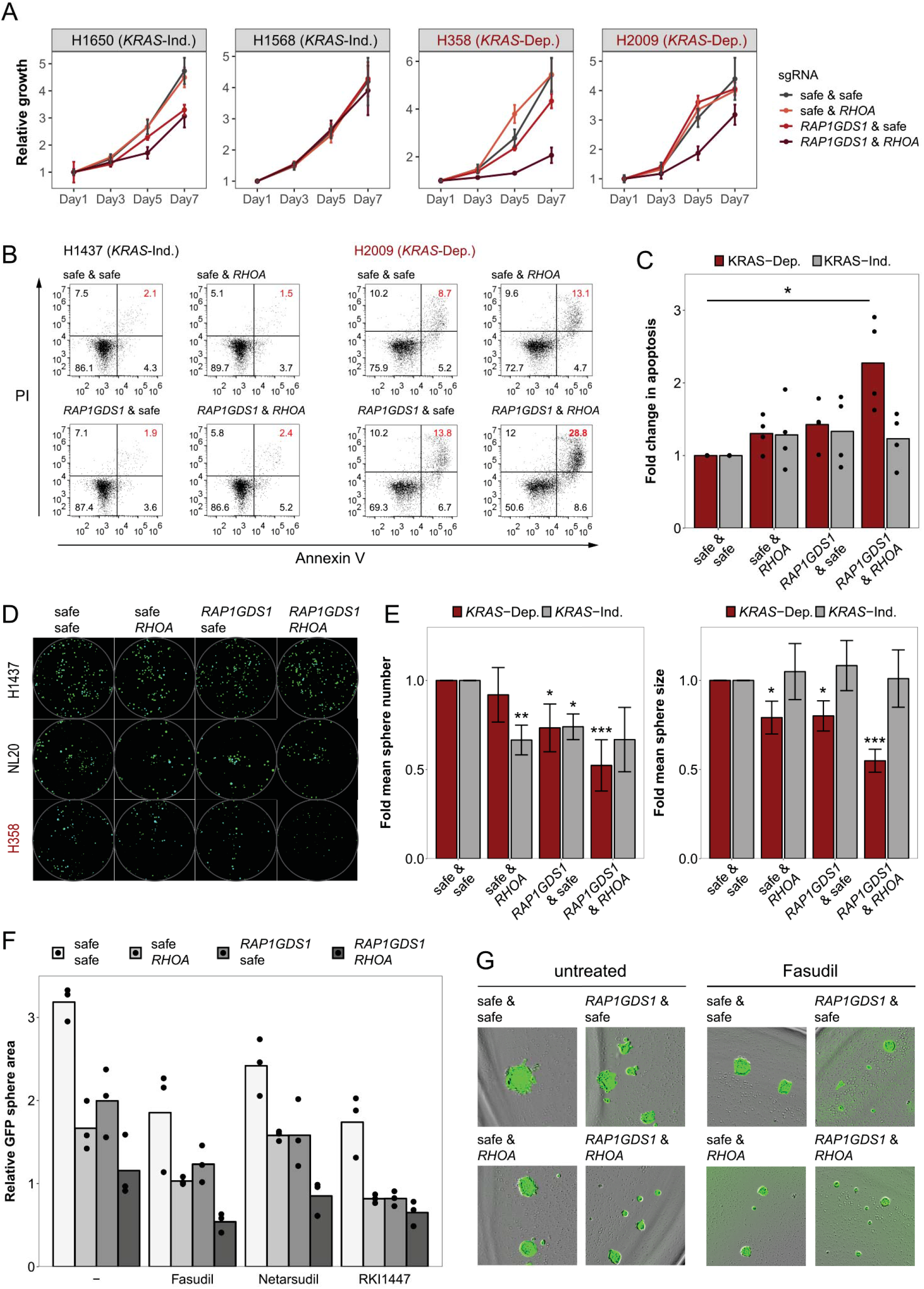
RAP1GDS1 *and* RHOA *form a Synthetic Lethal Gene Pair in RAS-Driven Lung Adenocarcinoma*. **A** Proliferation of two *KRAS*-independent (H1658, H1650) and -dependent cell lines (H358, H2009) infected with the indicated sgRNA pairs. Viability was measured on day 1, 3, 5 and 7 by Alamar blue. **B** Flow cytometry of *KRAS*-dependent (H2009) and -independent (H1437) cells by Annexin V-APC and propidium iodide. **C** Quantification of apoptosis. Red bars: *KRAS*-dependent cells (H2009, H358, H23); grey bars : *KRAS*-independent cells and HBECs (H1568, H1437, NL20). Bars represent means, * p<0.05 unpaired Student’s t-test. **D** Images of GFP-expressing spheroids in three lines expressing the indicated sgRNA pairs. **E** Quantification of sphere number (left) and size (right) in *KRAS*-dependent (H23, H358, H2009) and *KRAS*-independent (H1568, H1437, NL20) cell lines. Graph represents means of at least 3 independent experiments per cell line ± standard deviation, * p<0.05, ** p<0.01, *** p<0.001 unpaired Student’s t-test. **F** Spheroids of A549-Cas9 cells infected with the indicated sgRNAs treated with 10 μM Fasudil (ROCK inhibitor) or vehicle. **G** Quantification of sphere size in A549-Cas9 cells infected with the indicated sgRNAs and/or treated with ROCK inhibitors: Fasudil (10 μM) (as in F), Netarsudil (50 nM) or RKI1447 (500 nM).

Growth in spheroid culture reproduces important aspects of cancer biology not clearly modeled in 2D cultures (40). To further validate the importance of *RAP1GDS1/RHOA* synthetic lethality in *KRAS*-dependent cells, we tested the susceptibility of LUAD cell lines to dual *RAP1GDS1* and *RHOA* knockout in 3D culture. Individual knock-out of *RHOA* or *RAP1GDS1* had a minimal effect on sphere number and only a small effect on sphere size in *KRAS*-dependent cells (Figure 6D, E). However, double knockout significantly decreased both sphere number and sphere size in *KRAS*-dependent cells. In *KRAS*-independent cells, the loss of each of the genes individually lead to a decrease in sphere number, but the loss of both genes together did not synergize.

RHO Kinases are key effectors of RHOA function (41,42) and pharmacological inhibitors are available. We combined RHO-Kinase (ROCK) inhibition with *RAP1GDS1* knockout to determine if this would phenocopy the effect of the double knockout. Treatment of *KRAS*-dependent A549 cells with three different ROCK inhibitors (Fasudil, Netarsudil and RKI-1447) resulted in decreased sphere growth comparable to *RHOA* knockout alone (Figure 6F, G). However, ROCK inhibitors also showed effects in *RAP1GDS1/RHOA* double knockouts, suggesting that while RHO kinase is important for growth, it is not directly related to the synergy of *RAP1GDS1*/*RHOA* double knockout.

Although RAP1GDS1 might have general functions in membrane trafficking and cell growth, its effects in tumor cells appear to be surprisingly dependent on KRAS activation. Taken together, these studies identify a susceptibility of *KRAS* mutant LUAD to combined inhibition of both RAP1GDS1 and RHOA, uncovering RAP1GDS1 -- in its roles either as GEF or chaperone -- as a potential therapeutic target for treatment of *KRAS*-driven lung cancer.

## DISCUSSION

### The Complexities of RAS-Driven Cancer Motivate a Multiomic Mapping Approach

Genetic mutations cause pathway rewiring in cancer cells that both drives pathological behaviors and introduces susceptibilities that can be exploited in the clinic. Understanding this rewiring requires both knowledge of the individual molecules involved and the organization of interactions between them.

Large-scale tumor profiling efforts (1) and single gene CRISPR and RNAi screening efforts (9–11) have both made valuable progress in assembling a “parts list” of cancer pathways. However, these methods face major limitations identifying the linkage of cancer-related genes in cellular pathways. Cancer pathways have complex features that make potential pharmacological targets invisible to these methods. Such features identified in the RAS pathway, such as the RAF inhibitor paradox (43) and feedback inhibition (44) have important implications for cancer therapy development. The question of which pathway components are redundant is particularly important in RAS signaling, since many factors, including the RAS proteins themselves, are represented by multiple paralogs. Many agents in clinical use, like pan-RAF, pan-MEK, and pan-PI(3)K inhibitors, target classes of proteins rather than individual paralogs. Single-gene knockouts cannot mimic the family-wide inactivation caused by these drugs. Rewiring events may also introduce combination susceptibilities where cancer cells become sensitive to treatment by multiple agents. These possibilities, too, are not addressed by conventional screening methods.

We reasoned that we could address these shortcomings by systematic interaction mapping. By combining AP/MS and CRISPR-based CDKO genetic interaction (GI) mapping of RAS pathway components, we both expanded our knowledge of the RAS pathway and revealed new physical and genetic relationships between components. We have identified functional redundancies and differences between RAS, RAP, RAF, and RALGEF family members. The GIs we observe point to previously undescribed crosstalk between the RAF kinases and other RAS effector proteins. Finally, we have discovered new prospects for combination therapies involving both well-known targets like the RAF kinases or RAL GTPases and newer ones like RAP1GDS1. This combined PPI and GI map both aids our understanding of the RAS pathway and serves as a model for similar approaches to elucidate other disease pathways (Figure 7).

**Figure 7.**
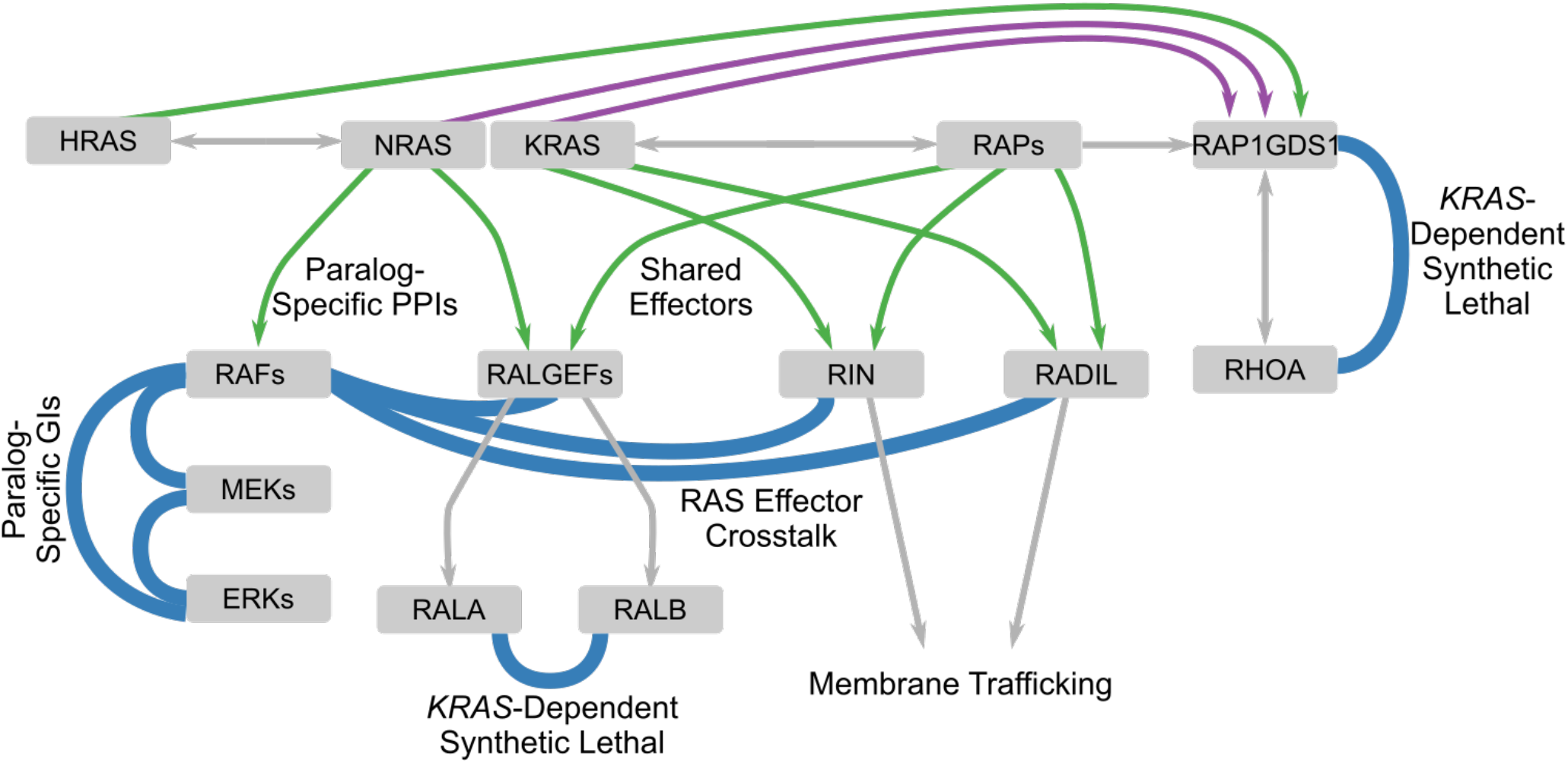
Genetic and Physical Interaction Mapping Reveal New Facets of RAS Pathway Biology. Schematic of RAS pathway features identified in this study. Among them are new, paralog-specific RAS/effector GI and PPIs, heterogeneity in GIs between MAPK pathway components, new roles in membrane dynamics for the RAS effectors RIN and RADIL, a striking intersection between RAS and RAP signaling, and synthetic lethal GIs between *RALA* and *RALB* and between *RAP1GDS1* and *RHOA* that are contingent on *KRAS* activity.

### Deep PPI Networks Illuminate New Facets of RAS Pathway Biology

Protein-protein interaction mapping by AP/MS has several features ideal for the particular challenges of RAS pathway biology. Among these are the unbiased identification of interacting proteins, the ability to capture PPIs as they occur *in situ*, and the ability to modulate sets of observed PPIs by judicious mutation of bait proteins.

To exemplify the utility of these experiments, we focused on two proteins that interacted with KRAS in its GTP-bound state: RIN1 and RADIL. While a RIN1-HRAS PPI was previously observed (20), RIN proteins have received little attention in RAS signaling. We demonstrate that RIN1 is a significant regulator of macropinocytosis, a process which has recently been specifically associated with RAS activation (Figure 2B,C)(22,23). These observations establish a mechanistic link between RAS and macropinocytosis and provide a context for more detailed screening (45).

We also demonstrate that KRAS binds and regulates RADIL via its RA domain similarly to known KRAS targets (Figure 2D-E, S2D-F). RADIL is a new RAS pathway member previously implicated in adhesion and cell migration (27,46), which we confirm. Though important for cell migration, further work is required to determine how RADIL aids small GTPase activity.

By combining the results of several AP/MS experiments in networks, we and other groups can make powerful comparisons among proteins, especially paralogs (16,47,48). By comparing the interactomes of H-, K-, and NRAS in A549 cells, we determined that the PPIs of HRAS with canonical RAS effector proteins are not observed in these cells. K- and NRAS, meanwhile, copurified many known RAS effector proteins, including the RAF kinase family and RALGEFs. Because the six RAS protein baits were expressed at comparable levels (Figure 1B), the strong differences likely reflect differences in biological regulation rather than artefacts of overexpression.

Virtually every RA-domain-containing protein observed interacting with any GTP-locked RAS protein *in vivo* co-purified with all GTP-locked RAS isoforms in cell-free systems. To judge by the PPI data, it appears that only a subset of these effectors are activated by a given RAS protein in a given cellular context. Therefore, some cellular regulation independent of the nucleotide state of each RAS protein modulates each RAS-effector PPI. Control of RAS and RAS effector localization is one likely mechanism by which different PPI patterns could be determined (49). Similarly, by mapping the different interactomes of three RALGEFs, we provide support for long-suspected differences in their cellular roles (29). By examining differences in PPIs between these different proteins, we can identify functional differences between similar proteins and more confidently map their links to different regulators, effectors, and cellular processes.

We were struck by the observation that proteins copurifying with RAS GTPases also copurify with RAP GTPases. Clustering analysis demonstrates that these two families share such strongly overlapping sets of interacting partners that they are not easily resolved (Figure 3C, D). While some links between these GTPases, and their sequence similarity, have been noted previously (32), the extent of PPI overlap suggests RAS-RAP crosstalk has hitherto been greatly underestimated. The regulation of RAP GTPase PPIs with RAS effectors, and their influences on RAS signaling, is an important problem.

Both our *in vitro* evidence and DepMap (11) suggest that many proteins linking these GTPases, like RIN1 and RADIL, are of measurable, but not of strong quantitative importance for cell line growth. However, the significance of macropinocytosis for RAS-transformed tumors is revealed free amino acids are scarce (22). RADIL, meanwhile, is required for the successful metastasis of MCF7 cells in mice (46). Therefore, the role of these effectors should be examined in other models that may better show their significance.

### The Combination of GI and PPI Mapping Produces Wiring and Rewiring Diagrams of Lung Adenocarcinoma

Having identified direct physical links between pathway components by PPI mapping, we conducted GI mapping to identify indirect functional links between components. The GI networks recovered predicted synthetic lethal GIs between the two MEK kinases (*MAP2K1* and *MAP2K2*) and the two ERK kinases (*MAPK1* and *MAPK3*). However, we found that GIs within the MAPK pathway differed between cell lines (Figure 4D, S6). Pan-RAF, pan-MEK and Pan-ERK inhibitors, while approved for some indications, often meet either toxicity limitations or adaptive resistance in clinical use (2,50). The variable and specific genetic GIs observed here support the investigation of isoform-specific RAF or ERK inhibition in RAS-driven cancers. Our subsequent comparison of GIs across multiple cell lines (Figure 5A) showed that MAPK pathway genes have more synthetic lethal GIs with non-MAPK genes in *KRAS*-dependent than -independent cells (Figure S5C). However, these GIs involve different gene pairs in different cell lines (Figure S7). The existence of these GIs, however, indicates a surprising degree of crosstalk between the MAPK pathway and other branches of the RAS pathway.

The heterogeneous susceptibilities we observe among cell lines agrees with (Shen et al 2017). However, the lines compared in that study (A549, HEK293, and HeLa) had very different primary sources, while the nine we compare are all LUADs. Though the similar provenance of these lines might be expected to yield similar GI patterns, we find few broadly consistent GIs (Figure 5E). These divergent GIs present a high-dimensional and somewhat daunting problem for the discovery of combination therapies. While PPIs help identify GIs, we observe most GIs between proteins distant in physical interaction space (Figure S5A) which could only be suggested by genetic or functional evidence. Notwithstanding, several GIs were sufficiently consistent to identify new predictions for combination therapies or the use of selective inhibitors.

For example, we found a strong synthetic lethal GI between *RALA* and *RALB* specific to *KRAS*-dependent lines (Figure 5B). While RALs are well-known RAS pathway members, they have yet to gain much interest as a therapeutic target. Though a RAL inhibitor has been developed targeting RAL interactions with RALBP1 (51), our PPI and GI data suggest RAL binding of the exocyst complex may be a more relevant endpoint.

Another noteworthy synthetic lethal GI between *KRAS* and *NRAS* was specific to cells *not* dependent on *KRAS* (Figure 5B). Given that the PPI results suggest that K-and NRAS both engage common effectors that HRAS does not (Figure 1C), we propose that by knocking out these two paralogs we have impaired the majority of RAS signaling. The fact that *KRAS*-dependent cells showed less synergy between *K*- and *NRAS* knockout suggests that these cells have indeed become specifically addicted to *KRAS*, rather than to RAS signaling more generally.

### RAP1GDS1 and RHOA as a Synthetic Lethal Susceptibility in RAS-Driven Lung Adenocarcinoma

The GI maps pointed to a KRAS-specific susceptibility of combined loss of *RAP1GDS1* and *RHOA*. Mutant RAS has previously been shown to require intact RHO signaling for transformation (52), and may be required for cell cycle progression in *KRAS* mutant cancers (39). RHO family members often harbor activating mutations in patient tumors (1).

RAP1GDS1 is a non-canonical GEF for several RHO family members. It promotes prenylation and membrane trafficking of RHO and RAS proteins, which are critical for their signaling (37,53). We identified RAP1GDS1 in the PPI data with all of the RAS and RAP GTPases as baits, making it a highly central member of this signaling network (Figure 1F, 3B, S1B, S2A). The importance of these binding partners in cancer signaling suggest RAP1GDS1 as a therapeutic target.

*In vitro* validation in 2D and 3D cultures of LUAD cell lines confirmed that dual *RAP1GDS1* and *RHOA* knockout selectively impaired growth of *KRAS*-dependent cells (Figure 6A-E,S8B), an effect we find tightly linked to induction of apoptosis. The extent to which this synthetic lethal GI derives from a two-hit insult to RHO signaling, or some other mechanism, requires further study. The fact that this GI manifests specifically in *KRAS*-dependent cells, however, offers new prospects for combination therapies.

### Conclusion

The concept of cancer pathway rewiring – a change of the linkage of cell biological pathways – underlies the core theories of targeted therapy development. In this work, we provide a survey of RAS pathway interactions in RAS-transformed LUAD. Unlike other recent cellular mapping approaches, in which the questions of interactions are neglected in favor of a wider survey of vulnerabilities or of mutational driving events, we map pathway wiring directly. In doing so, we have gained new insights into RAS-driven tumor biology and discovered new susceptibilities of RAS-driven tumors and provided a resource for other investigators to do the same.

## METHODS

### Plasmids

pMCB306, pKHH30 and p293 Cas9-BFP were previously described (12). pCMV-VSV-G and pCMV-dR8.2 dvpr were gifts from Prof. Bob Weinberg (Addgene plasmid #8454 and #8455) (54). pWPXLd LAP plasmids were also previously described(55). Cassettes were introduced in to GST-fusion, myc-fusion, and LAP-fusion plasmids by gateway cloning (Supplementary Methods). Point mutations in bait proteins were introduced by quickchange mutagenesis using primers in Supplementary Table 1.

### Cell Lines

HEK293, A549, H23, H358, H1437, H1568, H1650, H1975, and H2009 cells were obtained from American Type Culture Collection (ATCC). HEK293-LAP cell lines were generated from HEK293-FlpIn (55). Culture methods are described in Supplementary Methods.

### Affinity Purification / Mass Spectrometry

Affinity purification / mass spectrometry was carried out as described in (55), and quantified as in (16), with changes described in Supplementary Methods.

### Co-Immunoprecipitation Experiments

4×10^7^ cells of each line expressing a LAP-RAS mutant construct were lysed in 300 μL of 150mM KCl, 50mM HEPES pH 7.4, 1mM MgCl^2^ 10% glycerol, and 0.3% NP-40 (lysis buffer) supplemented with 3μL phosphatase inhibitor cocktail 2 (Sigma-Aldrich) and 1μg/mL each of leupeptin, pepstatin, and chymostatin (Sigma-Aldrich) on ice for 10 minutes. Cells were centrifuged for 15min at 4C. The concentration of each supernatant was measured by Bradford assay and back-diluted to 9mg/mL protein in 300μL lysate. For a 5% input sample, 9 μL were withheld and mixed with 4X LDS sample buffer +2.5% β-mercaptoethanol. 4.83 μL protein A beads conjugated to Rabbit αGFP antibody (Thermo Fisher) were added to the remaining supernatant. Samples were incubated 16h at 4C. Beads were washed 5 times with 200μL 200mM KCl, 50mM HEPES pH 7.4, 1mM MgCl^2^ 10% glycerol, and 0.3% NP-40 (lysis buffer), then immunoblotted as described(55), probed 16h at 4°C with rabbit αRAF1 (Abcam) and mouse αGFP (Invitrogen) at appropriate dilutions.

### Recombinant Protein Purification

Purifications of recombinant GST-tagged HRAS, KRAS, and NRAS with the indicated mutations performed as described in (55)). Briefly, proteins were purified from Rosetta 2 E. Coli cells transformed with plasmids encoding the indicated constructs.

### Cell-Free Co-Immunoprecipitation Experiments

Cell free co-immunoprecipitation experiments were performed as described in (55). Primary antibodies used were rabbit αGST (Sigma-Aldrich) and mouse αmyc clone 9E10 (EMD Millipore).

### Fluorescence Microscopy

Images were acquired on a Marianas spinning disk confocal (SDC) inverted microscope (Intelligent Imaging Innovations) using a Zeiss Plan-NEO 40x/1.3 Oil Ph3 objective, a Yokogawa csu22 confocal scanning unit and a Photometrics Cascade 1K camera.

### sgRNA Vectors

Small guide RNAs against human target genes or control safe-targeting sgRNAs were cloned into the MCB306 or KHH30 vectors (modified MCB306 with a human U6 promoter). For double sgRNA vectors the U6-sgRNA cassette was excised from the KH30 vector and cloned into the pMCB306 vector. All sgRNA sequences are listed in Supplementary Table 7. Lentivirus was produced in 293T cells as previously described (55) filtered, and applied directly to cells for infection at a MOI lower than 1. Incubation after infection is described in the relevant methods section. Clonal RIN1 knockout generation is described in Supplementary Methods.

### Macropinocytosis Assay

Macropinocytosis assays were conducted as described in (24), and elaborated in Supplementary Methods.

### Scratch-Wound Assay

Scratch-wound assays were performed as described (28), and are elaborated in supplementary methods.

### Quantitative Real-Time PCR

Quantitative Real-Time PCR for the RADIL knockdown experiments began with reverse transfection (Supplementary Methods), except that siRNAs were pooled. Cells were harvested and frozen 24, 48, and 72 hours post-transfection. Relative RNA levels were quantified using the TaqMan RADIL Assay (Thermo) with a compatible GAPDH Assay (Life) as a control probe using the TaqMan manufacturer’s protocol in 96-well MicroAmp Optical reaction plates (Applied Biosystems, N8010560)

### Clustering Analysis

Clustering analysis was conducted by the UPGMA method using the cosine distance metric, using the “linkage” method provided by the “scipy” module. Bait-prey interactions were binarized based on an FDR of < 0.05, and this adjacency matrix was used as input.

### CRISPR/CDKO screening

CDKO screening was conducted as previously described (Han et al. 2017), and is elaborated in Supplementary Methods.

### Colony formation assays

Cells were trypsinized, counted and 10,000 viable cells were seeded in complete medium into each well of a 6-well plate. After 10 days, colonies were stained with 0.2% methylene blue and quantified using ImageJ (NIH, Bethesda, Maryland, USA).

### Proliferation assays

Short term cell viability was measured using the Alamar Blue assay (Invitrogen). Cells were trypsinized and counted, 1,000 viable cells were seeded in 200ul of puromycin-containing medium into each well of a 96 well plate. Each condition was seeded in triplicate into one plate per time point. Alamar blue was added to the cells (20μL per well) 1, 3, 5 and 7 days after seeding, after which the plates were incubated for 4-6 h at 37C. Fluorescence was read using the Synergy Neo2 plate reader (BioTek, emission 555 nm/ excitation 596 nm).

### Sphere formation assay

For anchorage-independent sphere growth, cells were seeded into 24-well ultra-low attachment plates (20’000 viable cells per well) in 2ml of complete medium supplemented with 0.5% methylcellulose. The spheres were allowed to form for 9-20 days (depending on the cell line). GFP-positive spheres were imaged using the Leica DMi8 fluorescence microscope. Sphere size and number were quantified using ImageJ.

### Kinetic Growth Assays

Indicated Cas9-expressing A549, H23, H1437, and H1568 lines were infected with indicated lentiviral sgRNA constructs generated as described in Supplementary Methods. Media was replaced one day after infection. Cells were then incubated for two more days at 37C. Cells were then passaged into complete media culture medium plus 1μg/mL blasticidin and 2μg/μL puromycin (to select for the Cas9 cassette and sgRNA) and incubated two further days at 37C. Cells were then replated to 8×10^3^/mL in 24-well plates, with 6 wells per guide pair. 25 images were acquired in each well, and cell confluence measured, every two hours on an Incucyte S3 (Essen). Confluence data was fit to an exponential (base 2) equation, providing an estimate of doubling rate as plotted in Figure S8B.

### Statistical Analysis

Analyses of statistical significance were performed using the tests described in the texts, using the “scipy” library in Python. Where shown, 95% confidence intervals were calculated using the “statsmodels” library in Python (www.statsmodels.org).

Analysis of lesioning rates in Figure S7 relied on the provisional TCGA LUAD data from cbioportal.org. Using the EllipticalEnvelope method provided by scikit-learn, the mahalanobis distance of each gene from a common centroid based on relative mutation rates and CNA frequencies was calculated.

## ACKNOWLEDGEMENTS

We thank Alex Loktev, PhD and Kevin Wright, PhD and Christopher Adams, PhD and Ryan Leib, PhD for their helpwith mass spectrometry. We thank Dr. Ji Luo for advice on growth of lung cancer cell lines in spheroid culture.

## SUPPLEMENTARY METHODS

### Plasmids

The following plasmids were obtained from Harvard PlasmID database: pENTR RalGDS (HsCD00432064), pENTR RGL1 (HsCD00379137), pDONR221-RIN1 (HsCD00045692), pENTR201-RAP1B (HsCD00082137), pENTR201-RAP1A (HsCD00295731), pENTR221-RAP1B (HsCD00043785), and pENTR221 RAP1A (HsCD00296265). The following otherws were obtained from the National Cancer Institute Ras Initiative : pENTR255 Hs.ARAF (R207-E27), pENTR255 Hs.BRAF (R702-E23), pENTR255 Hs.RAF1 (R702-E25), pENTR255 Hs.KRAS4b (R750-E01), and pENTR255 Hs.KRAS4b G12D (R750-E05). The following plasmids were purchased from Genecopoeia : pDONR-RASGRF2 (GC-T8450), pDONR-RGL2 (GC-T1877-B), pDONR-RADIL (GC-Z3297), pDONR-RASIP1 (GC-Z1591). The following were purchased from Invitrogen: pENTR221 HRas (IOH39420), and pENTR 221 NRAS (IOH1768). Cassettes from the above donor vectors were cloned into destination cassettes by the Gateway method as described(1).

### RADIL-RA Subcloning and Mutagenesis

The RADIL RA domain was amplified from pCS2/6xMYC-RADIL using the primers 5’-GGGGACAAGTTTGTACAAAAAAGCAGGCTTAATGGACCCCGCCGAGCTCTCC-3’ and 5’-GGGGACCACTTTGTACAAGAAAGCTGGGTATTACTTCGCGCGACTCCGCTG-3’. The R99L mutation was introduced by quickchange PCR with the primers 5’-CCAGGGCGTACAGCTCCAGCGCC-3’ and 5’-GGCGCTGGAGCTGTACGCCCTGG-3’.

### Culture Method

Human embryonic kidney HEK293 cells were obtained from American Type Culture Collection (ATCC) and cultured in DMEM supplemented with 10% FBS, 2mM L-glutamine, 100 IU/mL of penicillin and 100μg/mL streptomycin, hereafter referred to as “complete DMEM”. cells and cultured in complete DMEM. A549 and H23 lung cancer cells were also obtained from ATCC and cultured in RPMI supplemented with 10% FBS, 2mM L-glutamine, 100 IU/mL of penicillin and 100μg/mL streptomycin, hereafter referred to as “complete RPMI”. Expression of LAP constructs in HEK293 cells was induced by doxycycline (1μg/mL treatment) for 24 hrs. prior to harvest for AP/MS experiments. A549 cells containing a doxycycline-inducible shKRAS construct, and all lung cancer cells containing Cas9 expression constructs were cultured in 2μg/mL puromycin. NL20 cells were cultured in Ham’s F12 medium (Gibco, #11765054) supplemented 2.7 g/L glucose (Sigma, #G8270), 1% penicillin–streptomycin-glutamine, 1x MEM Non-Essential Amino Acids (Gibco, #11140050), 1x ITSE (InVitria, #777ITS032), 10 ng/ml EGF (Humanzyme, #HZ-7012), 500 ng/ml hydrocortisone (Sigma, #H0888) and 4% FBS. BEAS-2B cells were grown in BEGM™ medium (Lonza, CC-2540B) on plates coated with 0.01 mg/mL fibronectin (Corning, #354008), 0.03 mg/mL bovine collagen type I (PureCol, #5005) and 0.01 mg/mL bovine serum albumin (Sigma, #A7030).

### Affinity Purification / Mass Spectrometry

Cells were cultured as previously described (2), harvested, and lysed. Bait and prey proteins were isolated from the lysate by tandem-affinity purification by a tandem-affinity purification process. The final eluate was fractionated by SDS-PAGE, and proteins were extracted from gel slices. These proteins were then prepared for LC-MS/MS and analyzed on an Orbitrap Fusion mass spectrometer (Thermo Fisher). MS/MS data was compared to an NCBI Genbank FASTA database containing all human proteomic isoforms with the exception of the tandem affinity bait construct sequence and common contaminant proteins. Spectral counts were assumed to have undergone fully specific proteolysis and allowing up to two missed cleavages per peptide. All data was filtered and presented at a 1% false discovery rate.

AP/MS data was quantified as described (3). Briefly, for individual genes whose products were identified in each AP/MS sample, we assigned a normalized spectral abundance factor (NSAF) to each gene (4). Using a panel of 116 other AP/MS experiments, we inferred expected lognormal background distributions for NSAFs of each gene using the median and scaled median absolute deviation for location and scale parameters, respectively. Each experiment was screened for correlation against each previous experiment, and highly correlated experiments (Pearson’s r>0.7) were removed from consideration prior to the inference of each gene-wise null distribution. For each gene-wise signal in an experimental dataset, we compare the signal of products of that gene to its background distribution, and it is from this comparison that *z*-scores and *p*-values are reported. We also conducted an experiment using renilla luciferase as a bait protein, since most negative control experiments were not conducted in A549 cells. This score allowed us to identify proteins more highly expressed in A549 cells relative to orthogonal controls. For most network diagrams (such as those in <u>NDex</u>, and in Figures S1 and S2), proteins enriched in the luciferase experiment (FDR<0.1) were removed from consideration. Data from the lluciferase experiment is included in Supplementary Table 2.

### RIN1 Knockout Generation

Clonal *RIN1*^−/−^ cells were generated as follows. Two different sgRNAs oligo duplexes were cloned by ligating of annealed oligo pairs 5’-TTGGCGTCTCCTTGGAGGGCTGTTTAAGAGC-3’ and 5’-TTAGCTCTTAAACAGCCCTCCAAGGAGACGCCAACAAG-3’ (sgRIN1-1) or 5’-TTGGAGCCTTGATGTTGAGGGGTTTAAGAGC-3’ and 5’-TTAGCTCTTAAACCCCTCAACATCAAGGCTCCAACAAG-3’ (sgRIN1-3). Meanwhile pMCB306 was digested with restriction enzymes BstXI and BlpI (New England Biosciences) per the manufacturer’s protocol. The linearized pMCB306 was ligated with each oligonucleotide duplex with T4 DNA ligase (New England Biosciences) per the manufacturer’s protocol and transfected into HEK293T cells (ATCC) using FuGENE6 (Fisher Scientific) alongside pCMV-dR8.2-dvpr and pCMV-VSV-G (Addgene). These cells were incubated at 37C for 24 hours, when the media was replaced. After 24 further hours, the supernatant containing assembled lentiviral particles was incubated with H23-Cas9 cells (see above). Two days after infection, this medium was replaced with culture medium plus 1μg/mL blasticidin (Fisher) scientific, then sorted on a Sony SH800 sorter into a 96-well plate (Corning) such that only one cell was present in each well. After expansion of these cells, a sample of each clonal population was harvested. Genomic DNA was purified using a DNAEasy Kit (Qiagen), and PCR-amplified with primers 5’-GTGGGAGGGAGAGGGATGG-3’ and 5’-CTGGAGGTCTCTGTGGACAC-3’ using Platinum Pfx DNA polymerase (Invitrogen) following the manufacturer’s protocol the cut sites of both guide RNAs are sufficiently close to each other that they could be both amplified and sequenced with the same primer pairs. The PCR products were then Sanger-sequenced by Sequetech (Mountain View, CA) using the primers 5’-AGGTCCTGAGCTGCCTGAG-3’ and 5’-TTGAAGCTTCTCTTGAATTTCTCC-3’. Sequences were analyzed by the ICE tool from Synthego (Menlo Park, CA). As plotted in Figure S3A, clone 1 is derived from sgRIN1-1 and clones 2 and 3 are derived from sgRIN1-3.

### Macropinocytosis Assay

Macropinocytosis was quantified as described (5). H23 cells were plated at 10^4^ cells/well in a 96-well optical dish (Eppendorf). 24 hours after replating, media was changed to complete RPMI + 10μM ARS853 (MedChemExpress) or equivalent volume DMSO (Sigma-Aldrich) and incubated at 37C. 24 hours later, media was replaced with serum-free RPMI + 10μM ARS853 or equivalent volume DMSO and incubated for 3 hours at 37C. Media was then replaced with the previous mixture, this time also containing 4μM CellTracker™ Green CMFDA (Thermo) and 0.1 mg/mL TMR-Dextran (Life Technologies). Cells were incubated a further 60 minutes at 37C. Cells were then washed 5 times with PBS + 0.5 mM MgCl^2+^ + 1mM CaCl^2+^ (PBS++) then fixed in PBS++ with 4% paraformaldehyde for 10 minutes at room temperature, then washed twice again in PBS++.

A maximum-intensity projection of green (λ^e×^=488) and red (λ^e×^=561) was calculated from 16 focal planes. Macropinocytosis was quantified by measuring the cellular area above an arbitrary but consistent green threshold and the area above an arbitrary consistent red threshold and dividing the red area over the green area, then normalizing each quotient to the mean of the control/vehicle condition. In figure 2C, bulk statistics are shown for the negative control and all three RIN1 knockout clones combined (n=81,71,74 for each clone).

### Scratch-Wound Assay

The scratch-wound assay was modified from a described protocol(6).2×10^5^ H23 cells were reverse-transfected with 30 picomoles siRNA. siRNAs were either a 1:1:1 mix of siRADIL −6, −7, and −8 or siGENOME Non-Targeting siRNA Pool #2 (“siCtrl”) (Dharmacon/GE). 48 hours post-transfection, cells detached using EDTA, back-diluted to 2.86×10^5^ cells/mL and plated in a 96-well ImageLock plate (Essen/Sartorius) previously coated with 1μg/cm2 bovine fibronectin (Sigma-Aldrich). After 8 hours, a scratch was made in each confluent monolayer and imaged every 30 minutes for 60 hours. Images were segmented using an in-house Python script using the scikit-learn and scikit-image libraries. Velocity was measured per pixel image from the second to penultimate timepoints. Each condition is represented by 5-6 scratches, but > 2×10^5^ separate velocity measurements.

### Construction of single and combinatorial CRISPR library (CDKO library)

For the single knockout CRISPR library that targets 223 genes derived from the PPI data, we first PCR-amplified total 3221 sgRNAs from a pooled oligo chip (Agilent). In this library, we included 1000 control sgRNAs (non-targeting sgRNAs with scrambled sgRNA sequences and Safe-sgRNAs that target non-functional regions of genomes) and 2221 sgRNAs to target 223 genes (10 sgRNAs / gene). The PCR amplified sgRNAs were then digested by BstXI/BlpI restriction enzymes and ligated into a MCB320 lentiviral vector which has mU6 promoter to drive expression of sgRNAs and EF1a promoter to drive expression of a puromycin-T2A-mCherry selection marker. For the CRISPR Double Knockout (CDKO) library that targets all possible combinations of the 119 genes selected from the single knockout CRISPR screening, we first PCR-amplified two sets of sgRNAs from a pooled oligo chip (Agilent). Each set contains identical sgRNAs with different cloning adapters. These sgRNAs contains 40 Safe-sgRNAs and 357 sgRNAs that target the 119 genes (3 sgRNAs / gene). For these 357 gene-targeting sgRNAs, we chose the best 3 sgRNAs out of 10 total sgRNAs per gene by their phenotypes in the single knockout CRISPR screening. We then digested one sgRNA set with BstXI and BlpI and ligated it into MCB320. We used Gibson assembly to ligate the other set of sgRNAs into KH30 lentiviral vector which has hU6 promoter to drive expression of sgRNAs. Lastly, hU6-sgRNA-tracrRNA cassettes in the KH30 library were cut out by XhoI and BamHI and ligated into the MCB320 library that was also cut with the same restriction enzyme pair. This generated the CDKO library which has 157,609 double-sgRNA combinations. The GI-directed sublibrary and Paralog sublibrary were constructed in the same way. The sgRNA set for GI directed sublibrary contains 6 Safe-sgRNAs and 60 sgRNAs to target the 20 selected hits. The set for Paralog sublibrary contains 8 Safe-sgRNAs and 72 sgRNAs to target 24 genes.

### CRISPR-Cas9 screens

For the CRISPR-Cas9 screens, we transduced the 9 LUAD lines with a lentiviral vector that contains EF1a-driven spCas9 fused to a blasticidin selective marker. Cell lines that constitutively express spCas9 were then selected with blasticidin. For the single knockout CRISPR screen that targets the 223 genes, we infected Cas9-expressing A549 cell line with the CRISPR library. For the 119 by 119 library CDKO library, A549-Cas9 and H23-Cas9 were used for the screen. The GI-directed sublibrary and Paralog sublibrary were transduced into all 9 LUAD lines. Once cells were transduced with CRISPR libraries, they were grown for 3 days, and selected with puromycin for another 3 days until the library pool became over 90% mCherry positive. Aliquots of the library were saved in liquid nitrogen as T0 samples, while the rest were used for the screen. All the screens were performed at ~1,000x cell number coverage for 21 days. Genomic DNAs from both T0 and Day 21 samples were then isolated and frequencies of sgRNAs were counted in the plasmid library, T0 genomic DNA samples, and Day 21 genomic DNA samples through deep sequencing. All screens were carried out in two experimental replicates and phenotypes of sgRNAs from replicates were normalized and pooled together to calculate final phenotypes and genetic interaction scores.

### Deep sequencing of sgRNAs

For each screen, genomic DNA was purified using QIAGEN Blood Maxi Kits from cells at ~1,000x coverage. Single or double-sgRNA cassettes were amplified with two rounds of PCRs utilizing Herculase II Fusion Polymerase (Agilent) as previously described (7) Briefly, 10 μg of genomic DNA was used as template in 100 μL PCR reaction for every ~5000 unique single/double-sgRNAs. The forward primers that bind to the 3’ end of the mU6 promoter and the reverse PCR primer that binds to the 3’ end of a typical single or double-sgRNA cassette were used in the first PCR. Illumina P5, P7, and 6-bp index were incorporated into the final PCR amplicons through the adapter primers during the second round of PCR. The PCR products were then gel-purified and sequenced on a NextSeq 550 (Illumina). Single-sgRNA cassettes were sequenced with single-read sequencing protocol with single index whereas doubles-sgRNA cassettes were sequenced with paired-end sequencing protocol with single index. Sequencing was performed at ~200x read-coverage for the single knockout CRISPR library and the 119 by 119 CDKO library and at ~1000x read-coverage for the GI-directed sublibrary and Paralog sublibrary.

### Measurements of phenotypes and genetic interaction scores

Growth phenotypes and genetic interaction scores (GI_T_ score) were calculated as previously described (7,8) In short, log2 fold enrichment of each single- or double-sgRNA between a plasmid library and a Day 21 sample is measured and adjusted by the median log2 fold enrichment of negative control sgRNAs (non-targeting sgRNAs and Safe-sgRNAs) included in a library. This value is then normalized by the standard deviation of all control sgRNAs included in the library to calculate a phenotype Z score (pZ). This normalization accounts for noise associated with each screen and would allow pooling of two replicates data. In the single CRISPR knockout library which has 10 sgRNAs per gene, growth phenotype of a gene is calculated as the median phenotype of corresponding 10 sgRNAs in one experimental replicate. In the double knockout library, phenotype of a gene pair is defined as the median phenotype of all existing 18 double-sgRNAs. In a typical CDKO library, there are 18 double-sgRNAs per gene pair since each gene has three sgRNAs and there are two possible orientations for a given gene pair (3 × 3 × 2 = 18). To calculate phenotypes of genes or gene pairs from two experimental replicates, normalized phenotypes of sgRNAs from two replicates are pooled together for a given gene (20 total pZ scores) or for a given gene pair (36 total pZ scores) and the median phenotype of pooled sgRNAs are used as final phenotypes of a gene or a gene pair. In the single knockout CRISPR screen, we performed Mann-Whitney U (MWU) test between the distribution of all single sgRNAs targeting a gene and that of all negative control sgRNAs in a library to calculate MWU p-value of the gene. False discovery rates (FDRs) of genes are then estimated from these MWU p-values using Benjamini-Hochberg procedure. We used GI_T_ score as a statistical score of genetic interactions to account for both strength and consistency of genetic interactions among double-sgRNAs targeting a single gene pair. Briefly, we first calculated an expected phenotype of a double-sgRNA by summing single phenotypes of two sgRNAs in the pair. We subtracted this expected phenotype from observed phenotype of the double-sgRNA to calculate genetic interaction of the double-sgRNA. The sign of genetic interaction is re-defined in this step as follows. If observed growth phenotype is more toxic than the expected phenotype, negative sign is assigned to the genetic interaction. If observed phenotype is less toxic than the expected, then positive sign is assigned. To calculate genetic interaction of a given gene pair, the median genetic interactions of corresponding double-sgRNAs is calculated and that of non-interacting double-sgRNAs (any double-sgRNAs that contain at least one Safe-sgRNA) is subtracted from the value. This adjusted value is then normalized by the noise levels associated with the double-sgRNAs targeting the gene pair and the non-interacting double-sgRNAs to calculate the GI_T_ score of the gene pair.

### Network Diagrams

Network diagrams were rendered in Cytoscape (9).

## SUPPLEMENTARY TABLES

*Supplementary Table 1*

Related to Figure 1

Primers for quickchange mutagenesis of the indicated bait proteins used in AP/MS experiments.

*Supplementary Table 2*

Related to Figure 1

This table lists genes whose protein products were identified by AP/MS experiments in HEK293 cells or A549 cells on the indicated sheets. The columns are as follows: Bait, gene of bait protein; Prey, gene identified of prey protein; NSAF : The log_10_ transform of the Normalized Spectral Abundance Factor (see Methods), with greater numbers indicating stronger signals; Background Values : NSAFs for other experiments, used to infer the background distributions as described above.

*Supplementary Table 3*

Related to Figure 1

This spreadsheet contains 6 tabs, with each tab corresponding to a cell line and GO namespace. For each indicated bait (columns), all hits detected with FDR ≤ 0.05 were searched for enrichment for the indicated term (row) using a background set of all protein-coding genes for which at least one GO term existed by Fisher’s Exact Test. The values of the table are the FDR-corrected p-values from that test by the Benjamini-Hochberg method.

*Supplementary Table 4*

Related to Figure 4

Results from the single-target sgRNA screen in A549-Cas9 cells. The columns are as follows : Symbol : HUGO gene symbol; EnsemblID : EnsEMBL identifier; GeneID : Entrez Gene ID; GeneInfo : Name of protein encoded by the gene; Localization : GO cellular localization terms; Process : GO cellular process terms; Function : GO molecular function terms; #Element: Number of identified sgRNAs in sequencing; Phenotype (pZ) : median log_2_-fold enrichment of sgRNAS targeting this gene over the course of the experiment; Tscore : GI_T_ score; MWU P-Val; p-value of gene depletion or enrichment by Mann-Whitney U test relative to Safe-sgRNAS; MWU Adjusted Pval : FDR as determined by the Benjamini-Hochberg method.

*Supplementary Table 5*

Related to Figure 4

Results from the first pairwise sgRNA screen in A549-Cas9 and H23-Cas9 cells. Columns are as in the previous table, except that they refer to Gene A or Gene B as indicated, where the symbol for GeneA comes alphabetically before the symbol of Gene B. Test statistics reflect comparisons with inferred expected phenotypes from Gene A/safe or Gene B/safe sgRNA pairs (see Methods).

*Supplementary Table 6*

Related to Figure 5

Results from the “GI-Directed” sublibrary screen. Different metrics are reported on each of the three tabs of the spreadsheet as indicated. Columns correspond to individual replicates in the indicated cell line.

*Supplementary Table 7*

Related to Figure 5

Results from the “Paralog” sublibrary screen. Different metrics are reported on each of the three tabs of the spreadsheet as indicated. Columns correspond to individual replicates in the indicated cell line.

*Supplementary Table 8*

Related to Figures 4 and 5

Table of sgRNA sequences.

## SUPPLEMENTARY FIGURE LEGENDS

**Supplementary Figure S1.**
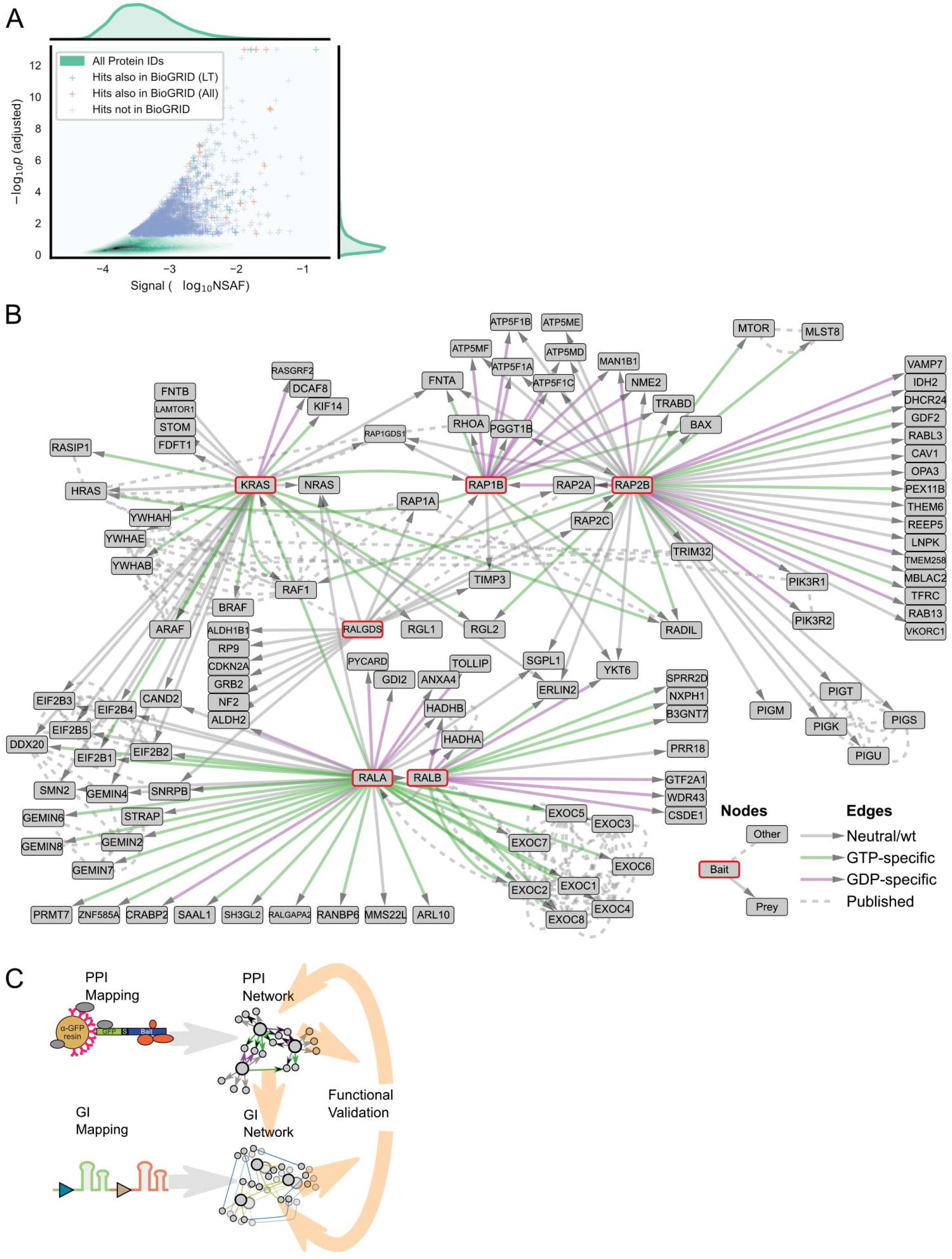
**Related to Figure 1 A** Volcano contour plot of gene-wise protein identification across all AP/MS experiments. Hits (FDR<0.05) are plotted as crosses, with hits also identified in BioGRID by low-throughput (LT) methods in green, by any method in orange, and hits not identified in BioGRID in blue. **B** Network diagram showing curated PPIs from AP/MS experiments conducted in HEK293 cells. Green lines indicate interactions specific to the GTP-locked mutant, while purple lines indicate interactions specific to the GDP-locked mutant for GTP-locked baits. PPIs with no nucleotide bias, or from non-GTPase prey proteins (i.e. RALGDS) are in grey. Dashed lines indicate interactions derived from BioGRID or HuMAP. Mutant and bait proteins are displayed using their source gene name. Corresponding data can be found in Supplementary Table 1 or on <u>NDEX</u>. **C** Schematic illustrating a generalizable version of the approach followed by this paper. PPI mapping is conducted to produce a physical interaction map, whose members are then used as targets for genetic interaction mapping. After appropriate follow-up experiments, each procedure can be repeated for previously unexplored parts of the earlier networks.

**Supplementary Figure S2.**
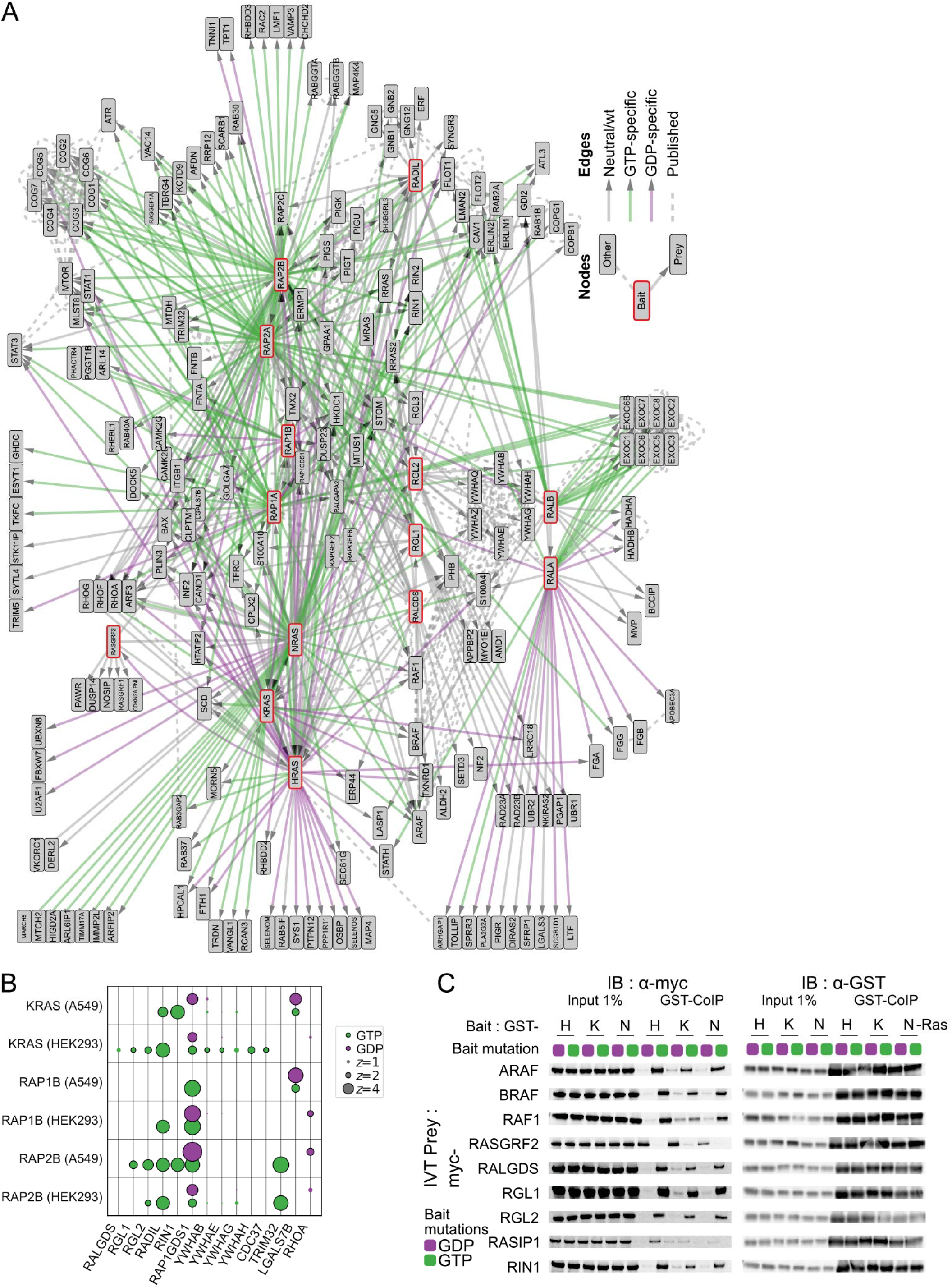
**Related to Figure 1 A** Network diagram showing curated PPIs from AP/MS experiments conducted in A549 cells, plotted as in S1A. **B** Bubble plot of GTPases identified co-precipitating with RAS and RAP GTPases in A549 cells. **C** Immunoblots of *in vitro* co-immunoprecipitations of RAS proteins with their putative effectors. Myc-tagged prey proteins were expressed in wheat germ extract and incubated with purified recombinant GST-tagged prey proteins with the indicated mutation. This data includes that shown in Figure 1E and loading controls.

**Supplementary Figure S3.**
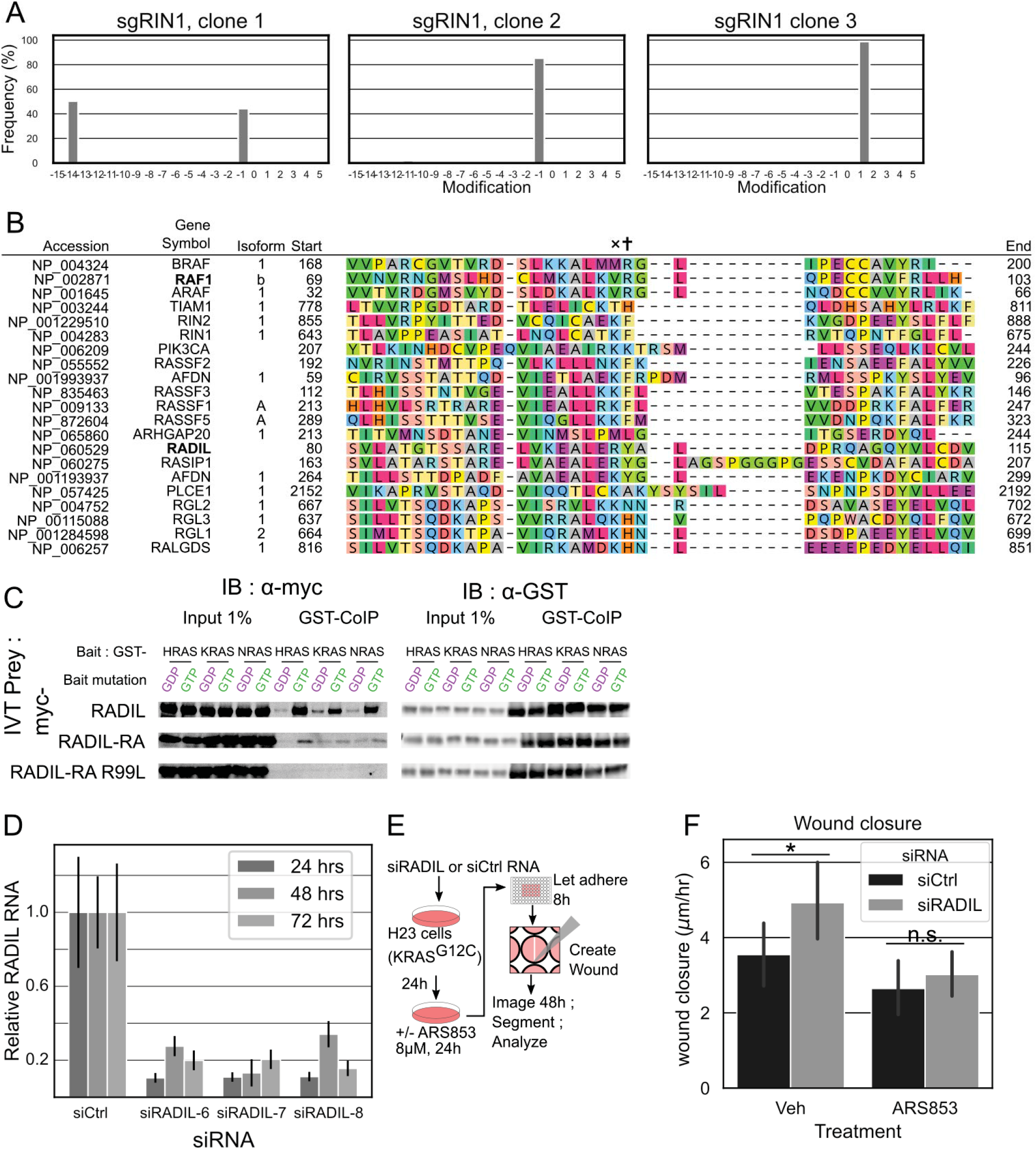
**Related to Figure 2 A** Alteration plot showing indel frequencies in 3 RIN1 KO clones of H23 cells used for Figure 2B. The population arising from clone 1 has one allele with a 14bp deletion and a 1bp deletion in the second; cells from clone 2 have a 1bp deletion in both alleles, and cells from clone 3 have a 1bp insertion in both alleles. Plot was generated from analysis using the ICE tool from synthego.com. **B** Sequence alignment of selected proteins containing RAS-association domains. The arginine corresponding to RAF1 R99 is marked with †; that corresponding to RADIL R89 with ×. **C** Co-immunoprecipitation of indicated myc-tagged constructs with recombinant mutant GST-tagged H, K, and NRAS bait proteins. **D** Quantitative Real-Time PCR of 3 indicated siRNAs against RADIL (see supplementary methods). Total RNA was isolated from transfected H23 cells at 24, 48, and 72hrs post-transfection and the relative levels of RADIL transcript were quantified. For experiments with RADIL knockdown, a 1:1:1 pool of three siRNAs were used. *n=4* for all samples. **E** Schematic of scratch-wound assay. Cells transfected with anti-RADIL siRNA or a negative control were replated into a confluent monolayer. A wound was introduced in the monolayer, and the closure of the wound was monitored over time. **F** Quantification of cellular velocity into the wound over 24 hours; RADIL knockdown increased migration speed, but not when KRAS was inhibited. 5-6 wounds were quantified per condition, imaged hourly (Supplementary Methods). * : *p*<0.05 by Welch’s *t*-test.

**Supplementary Figure S4.**
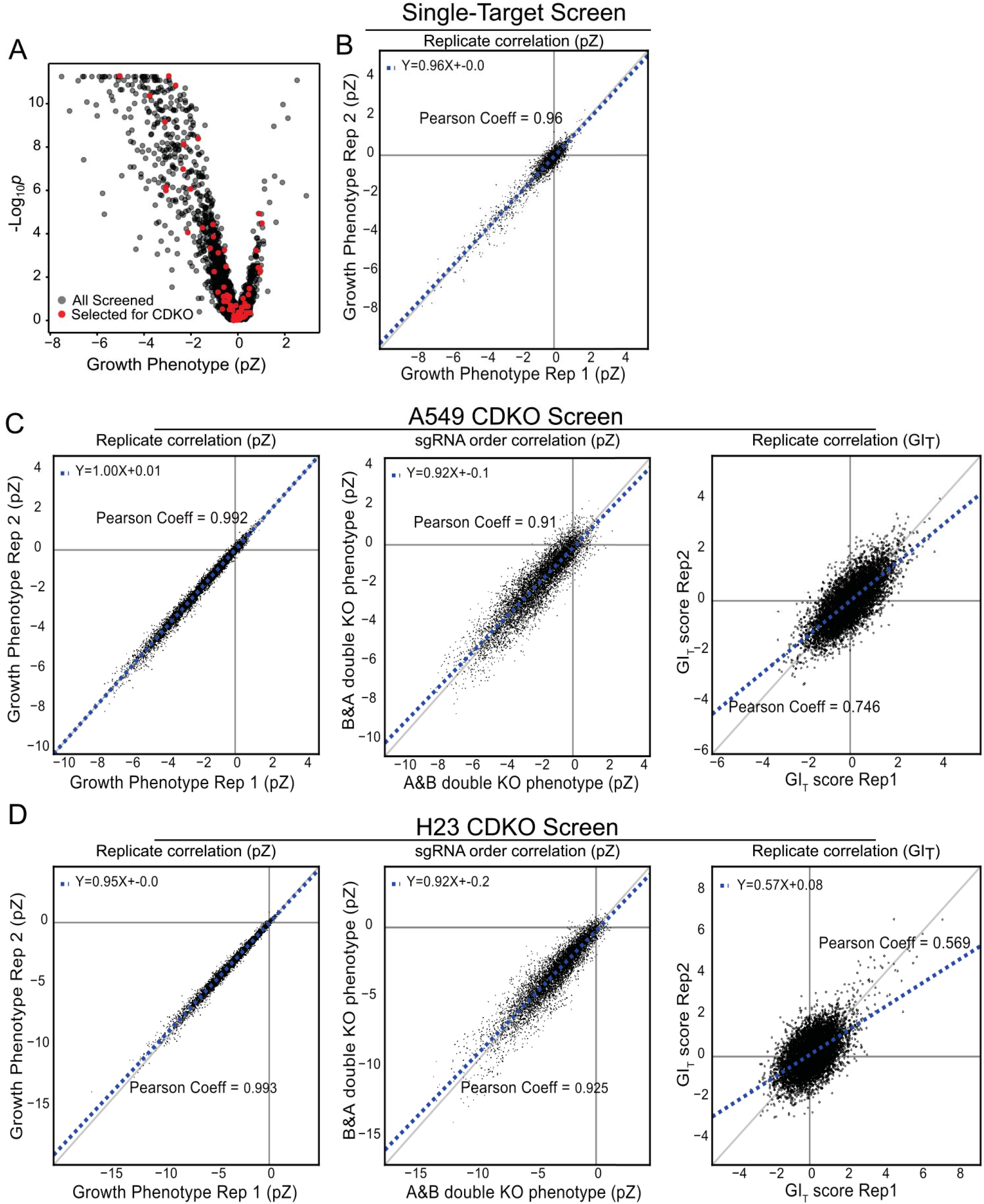
**Related to Figure 4 A** Volcano plot showing effects of gene knockouts in the single sgRNA screen in A549 cells. Points marked in red correspond to genes chosen for the large pairwise library; those with too deleterious a growth phenotype were excluded, and others were prioritized on the basis of their place in the network. **B** Growth phenotypes were highly reproducible between experimental replicates in the single sgRNA screen in A549 cells. **C** Double knockout growth phenotypes measured in the large CDKO screen in A549 cells were highly reproducible between experimental replicates (left panel). In the screen, double knockout phenotypes of gene pairs were compared between two different orientations (A&B, B&A) and showed minimal positional bias (middle panel). Genetic interaction scores were highly reproducible between experimental replicates (right panel) **D** As in A549, the large CDKO screen in H23 also showed highly reproducible growth phenotypes (left panel) and genetic interaction scores (right panel) between experimental replicates and also showed minimal positional bias (middle panel) for double knockout phenotypes of gene pairs.

**Supplementary Figure S5.**
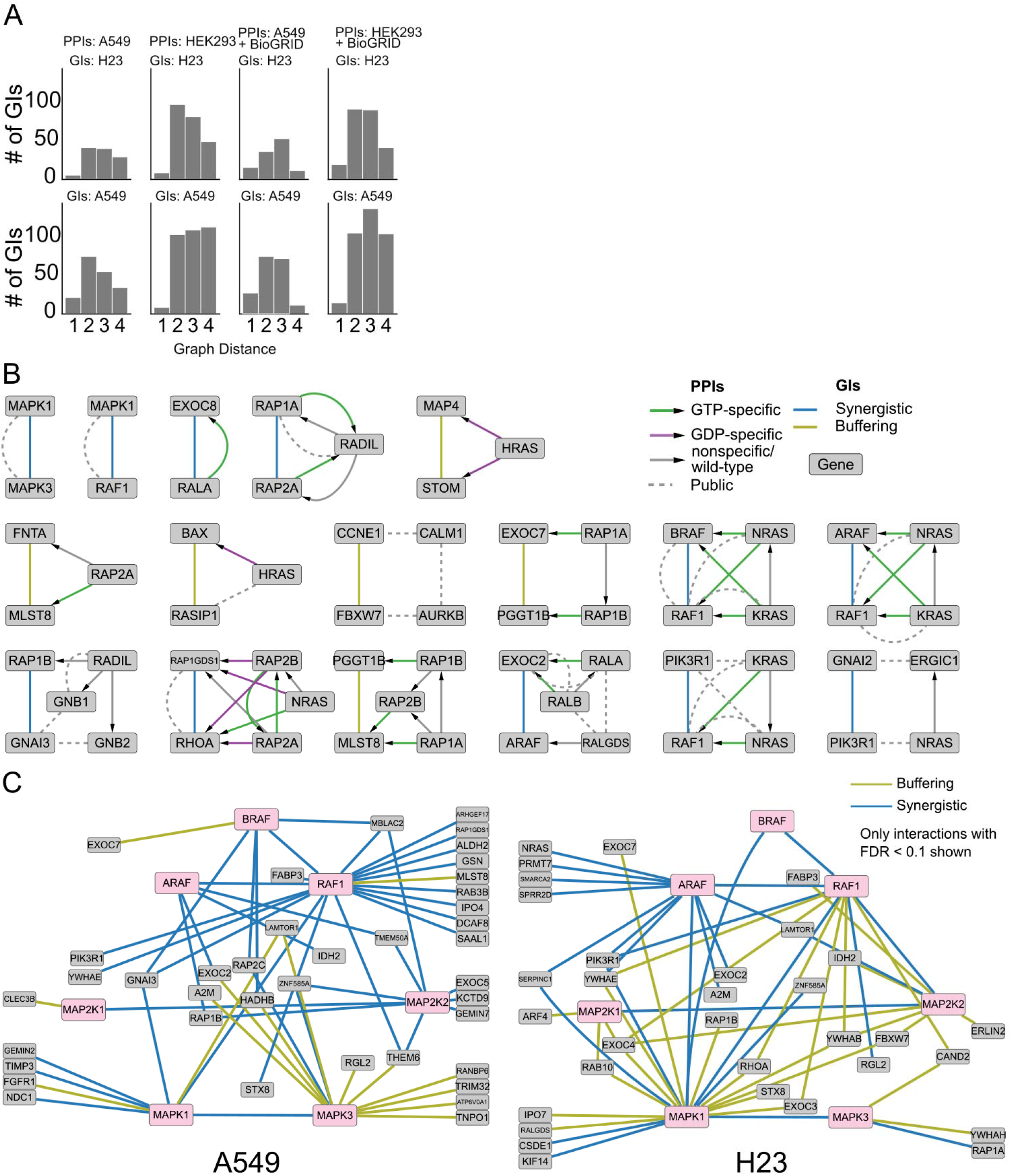
**Related to Figure 4 A** Histograms showing the graph distance (in a PPI network) between pairs of genetically interacting genes. The PPI networks were assembled from either the HEK293 data or the A549 data. In the ‘+BioGRID’ histograms, the indicated data was supplemented with public PPI from BioGRID. **B** Subgraphs of protein-protein (PPI) and genetic (GI) interaction data. Each subgraph contains one genetic interaction observed to be consistent in A549 and H23 cells, and the shortest physical interaction path that joins both interactors. Paths were determined from in-house PPI data as well as published data from BioGRID. **C** Network diagram of GIs in H23 and A549 cells from the large library. Only genetic interactions involving at least one MAPK pathway component (pink) are shown.

**Supplementary Figure S6.**
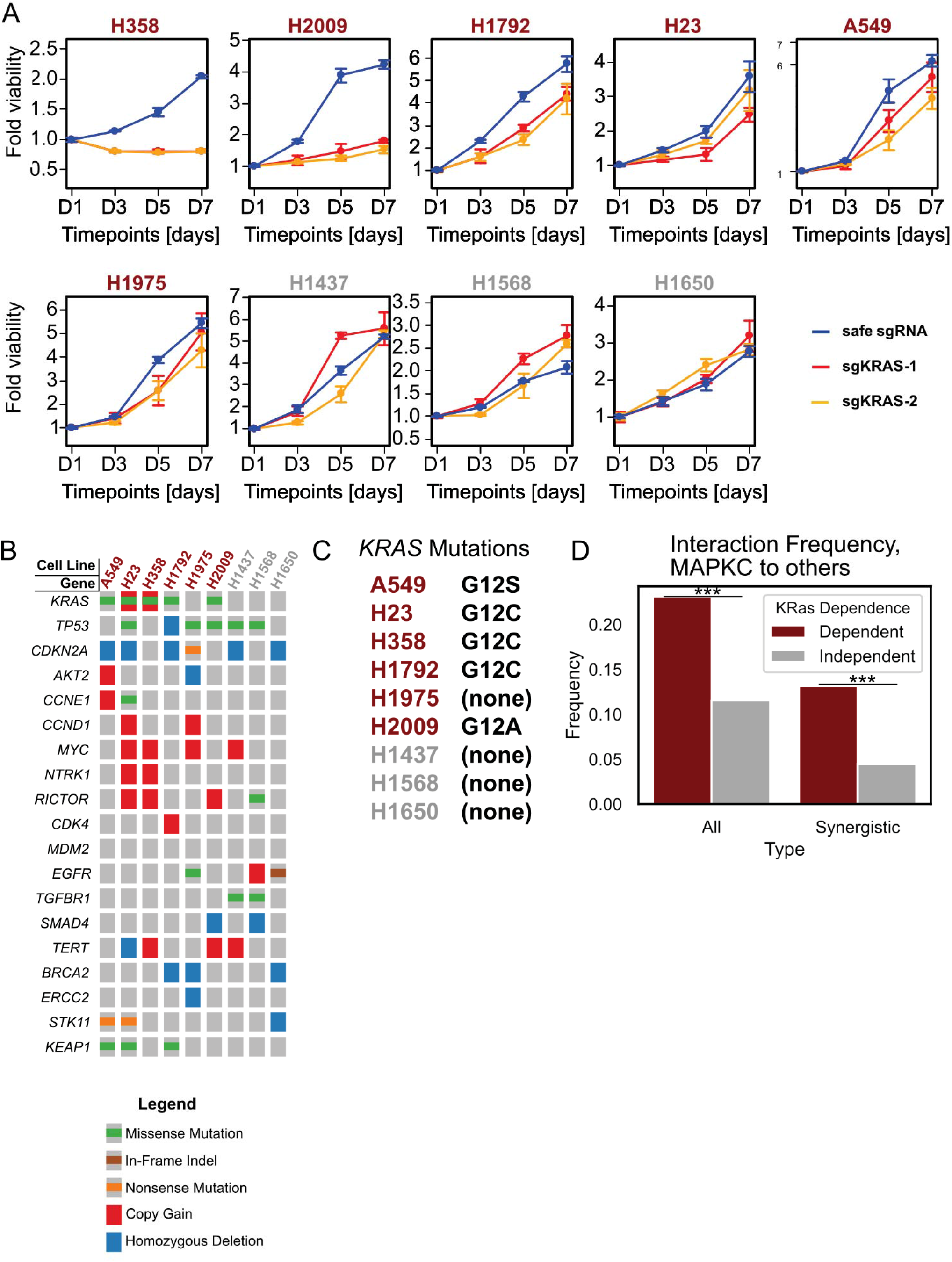
**Related to Figure 5 A** Proliferation assay of the indicated cell lines across 4 timepoints post seeding after transfection with the indicated guides targeting *KRAS*. **B** List of KRAS protein mutations in the indicated cell lines. **C** Oncoprint of selected mutations in the indicated cell lines from the Cancer Cell Line Encyclopedia. The selected cell lines are heterogeneous in terms of non-KRAS genetic alterations. **D** Frequency of genetic interactions (observed interactions /possible interactions) where one and only one partner is a MAPK pathway component (ARAF, BRAF, RAF1, MAP2K2, MAP2K4, MAPK1, MAPK3). Interactions involving these proteins are much more common in *KRAS*-dependent cancers. *** p < 0.001 by Fisher’s Exact Test.

**Supplementary Figure S7.**
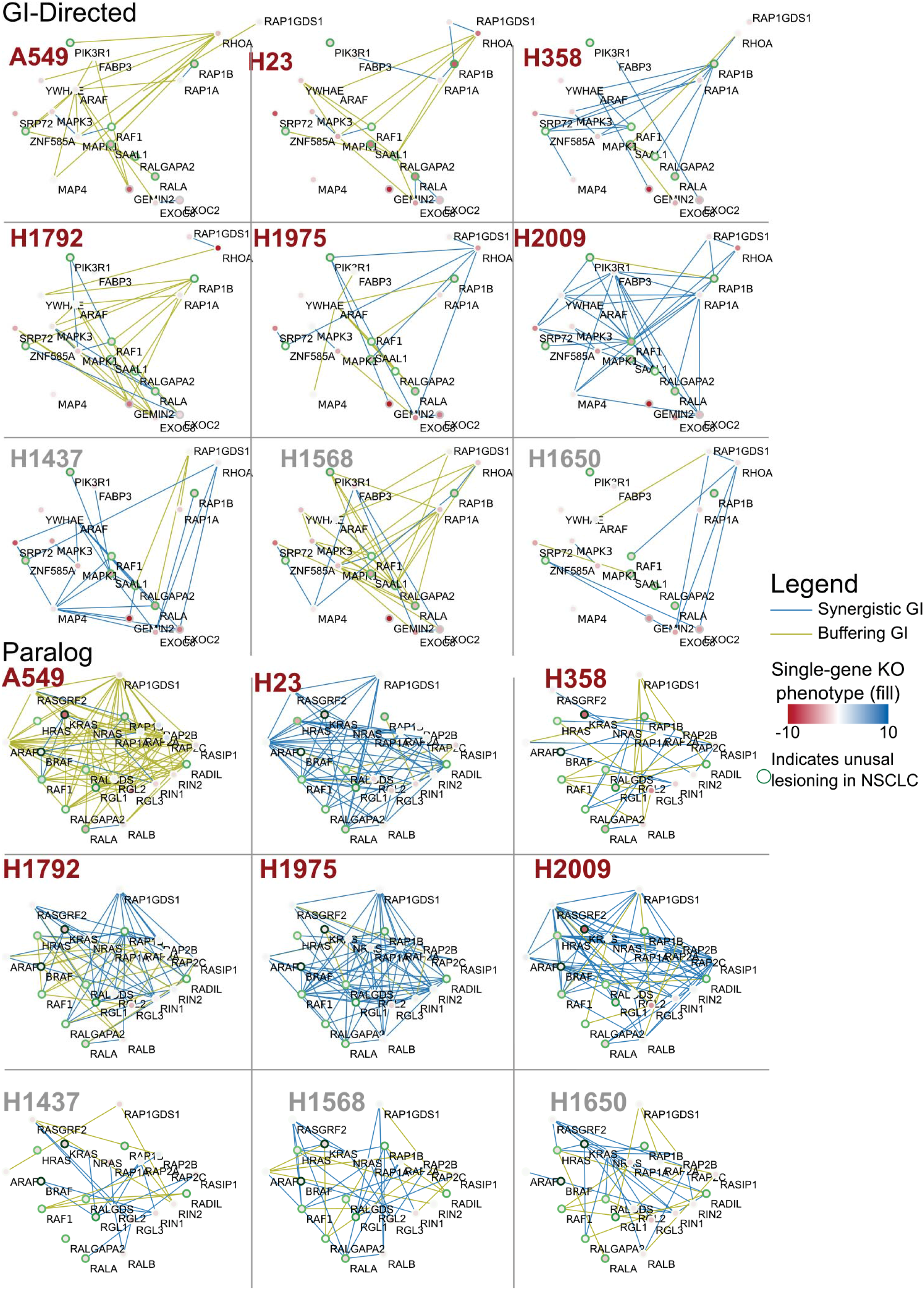
**Related to Figure 5** Complete set of all genetic interactions with FDR < 0.1 from all lines with data from both the “GI-directed” and “Paralog” libraries. Node (gene) locations are analogous between all networks. The node fill color is indicative of the single-gene knockout phenotype in that experiment; the green border indicates unusually frequent lesioning in NSCLC (see STAR methods).

**Supplementary Figure S8.**
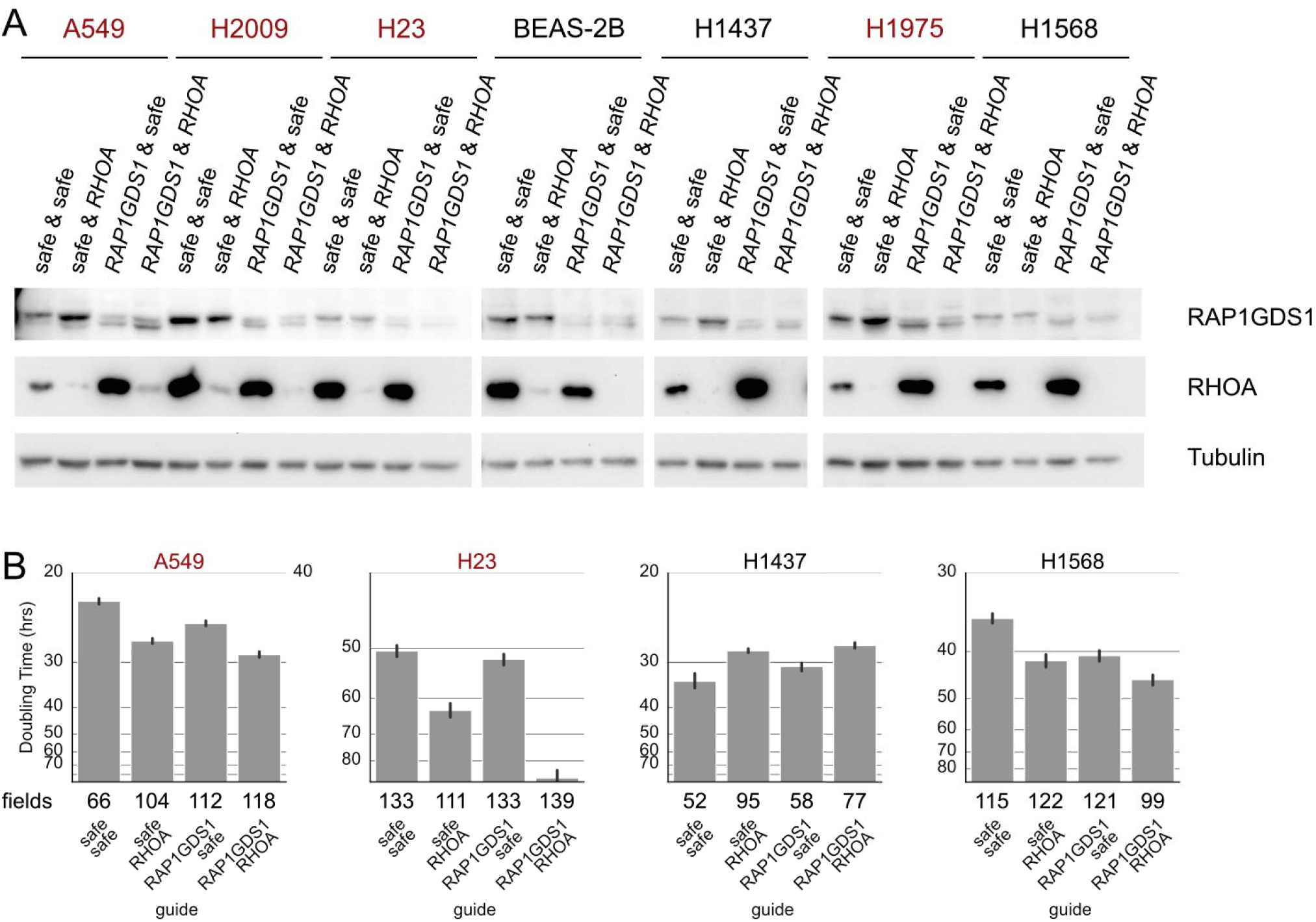
**Related to Figure 6 A** Lysates of Cas9-expressing cells infected with the indicated double sgRNA vectors were analyzed by immunoblotting for indicated proteins. **B** Doubling times of indicated Cas9-expressing cell lines infected with indicated sgRNA pairs inferred from real-time growth assays.

